# Spatiotemporal NF-κB dynamics encodes the position, amplitude and duration of local immune inputs

**DOI:** 10.1101/2021.11.30.470463

**Authors:** Minjun Son, Tino Frank, Thomas Holst-Hansen, Andrew Wang, Michael Junkin, Sara S Kashaf, Ala Trusina, Savaş Tay

## Abstract

Infected cells communicate through secreted signaling molecules like cytokines, which inform nearby cells about the type, severity and location of pathogens. How differences in cytokine secretion affect inflammatory signaling over space and time, and how responding cells decode information from propagating cytokine signals are not understood. By computationally and experimentally studying NF-κB dynamics in co-cultures of signal sending cells (macrophages) and receiving cells (fibroblasts), we found that cytokine signals are transmitted by wave-like propagation of NF-κB activity and create well-defined cellular activation zones in a responding cell population. Remarkably, NF-κB dynamics in responding cells can simultaneously encode information about cytokine dose, duration, and distance to the cytokine source. Spatially-resolved transcriptional analysis revealed that responding cells transmit local cytokine information to distance specific pro-inflammatory gene expression patterns, creating “gene expression zones” in the population. Despite single-cell variability, the size and duration of the signaling zone is tightly controlled by the macrophage cytokine secretion profile. Our results highlight how macrophages tune their cytokine secretion dynamics to control signal transmission distance, and how NF-κB interprets these signals to coordinate inflammatory response in space and time.

## Introduction

Response to infection requires cellular coordination in a wide range of temporal and spatial scales. Innate immune response starts with pathogen detection by sentinel cells like macrophages, which coordinate gene expression in nearby cells through secreted cytokines. Propagating cytokine signals relay information about the type, severity, timing and location of pathogen inputs. Signaling pathways like NF-κB play a vital role in interpreting such signals (Hoffmann and Baltimore, 2006). Signals from self (cytokines or chemokines) as well as from pathogens (bacterial or viral components) both activate NF-κB transcription factors, which rapidly shuttle into the cell nucleus and lead to expression of signal specific response genes.

Previous studies showed that temporal characteristics of NF-κB is key for immune regulation, and dysfunction of NF-κB dynamics is associated with diverse pathologies from autoimmune disorders to cancer (Lawrence, 2009; Pereira and Oakley, 2008). Extensive study of single-cell NF-κB dynamics resulted in improved mechanistic insight in pathway activation and control of signal-specific gene expression, and led to the development of accurate computational models of immune gene regulation (Cheng et al., 2021; Hoffmann et al., 2002; Kellogg and Tay, 2015; Nelson et al., 2004; Tay et al., 2010). However, NF-κB signaling in a spatial context has not been studied in detail, and the cellular and molecular mechanisms behind spatial coordination of cells during innate immune response are not clear.

Immune cells must coordinate local and global tissue-level responses at various spatial scales (Altan-Bonnet and Mukherjee, 2019; Voisinne et al., 2015). Theoretical studies predicted that cytokine signals create a sharply-defined cell activation zone in tissue (Bagnall et al., 2018; Cheong et al., 2006), while a recent *in vivo* study showed that size of the signaling range depends on the density of cytokine-consuming cells (Oyler-Yaniv et al., 2017). Despite these earlier works, studies addressing spatio-temporal aspects of signaling are rare, mainly due to the difficulties in tracking dynamic processes between cells. Quantitative understanding of spatio-temporal cellular communication could lead to broadly applicable models on the emergence and progression of infection, immunity, and autoimmunity, and can lead to developing effective therapies (Chen et al., 2017; Thaiss et al., 2016; Xu et al., 2019).

Several questions on spatial regulation of immune signaling are outstanding. First, cellular responses are naturally variable and subject to molecular noise. For example, cytokine secretion from individual macrophages show different concentrations and temporal changes under identical pathogen inputs (Chen et al., 2019; Junkin et al., 2016; Xue et al., 2015). Furthermore, signal receiving cells like fibroblasts respond to cytokines in a highly variable manner (Adamson et al., 2016; Adelaja et al., 2021; Tay et al., 2010). How single-cell variability impacts information transfer, signaling range, and gene expression at various distances from the infection site are not understood.

Additionally, it is not known if signaling networks can determine distance and location of the signal source in tissue. The dose and duration of a cytokine signal provides information about the severity and persistence of the pathogen load in the tissue microenvironment. Secreted cytokines form concentration gradients, which could enable responding cells to locate the pathogen input. However, simultaneously determining the dose and location of a cytokine input is challenging because high dose in gradient could indicate either proximity to the source, or a high initial cytokine release. Thus, measurement of the dose alone may not provide enough information to the cells about the severity and location of the pathogen input.

While it is not clear if signaling networks are capable of simultaneously decoding pathogen load and location, dynamical characteristics of NF-κB provide potential for such multi-dimensional information processing (Cheong et al., 2011; Selimkhanov et al., 2014). Time-course of a local cytokine signal depends on initial cytokine release and distance, and NF-κB dynamics in single cells is highly sensitive to such changes (Son et al., 2021). This suggests the intriguing possibility that NF-κB may simultaneously decode the amplitude, duration and location of a pathogen input. Determining spatio-temporal processing abilities of signaling pathways would greatly improve our understanding of how immune cells spatially coordinate their inflammatory responses.

To study how localized cytokine signals control NF-κB signaling and gene expression in space and time, we developed a microfluidic co-culture system that mimics conditions during inflammatory signaling after infection (Figure 1C). Using this system and mathematical modeling, we investigated how pathogen detection by macrophages is broadcast to nearby cells, and how these signals are interpreted by the NF-κB network in individual cells at different locations. Through computational simulations and live-cell analysis, we found that the signaling range and duration are highly variable, and depend strictly on the rate of cytokine production by macrophages. Surprisingly, we find that NF-κB network can simultaneously measure the dose, duration and distance of a cytokine signal, despite the variability in the signal receiving cell population. Spatially-resolved gene expression measurements showed that cells convert spatially varying signals into distinct spatial gene expression patterns within the responding cell population, creating “gene expression zones”. Our studies resulted in a computational model of NF-κB signaling in space and time and revealed how cells use NF-κB dynamics to extract complex information from their environments and coordinate their responses to pathogen inputs.

**Figure 1.**
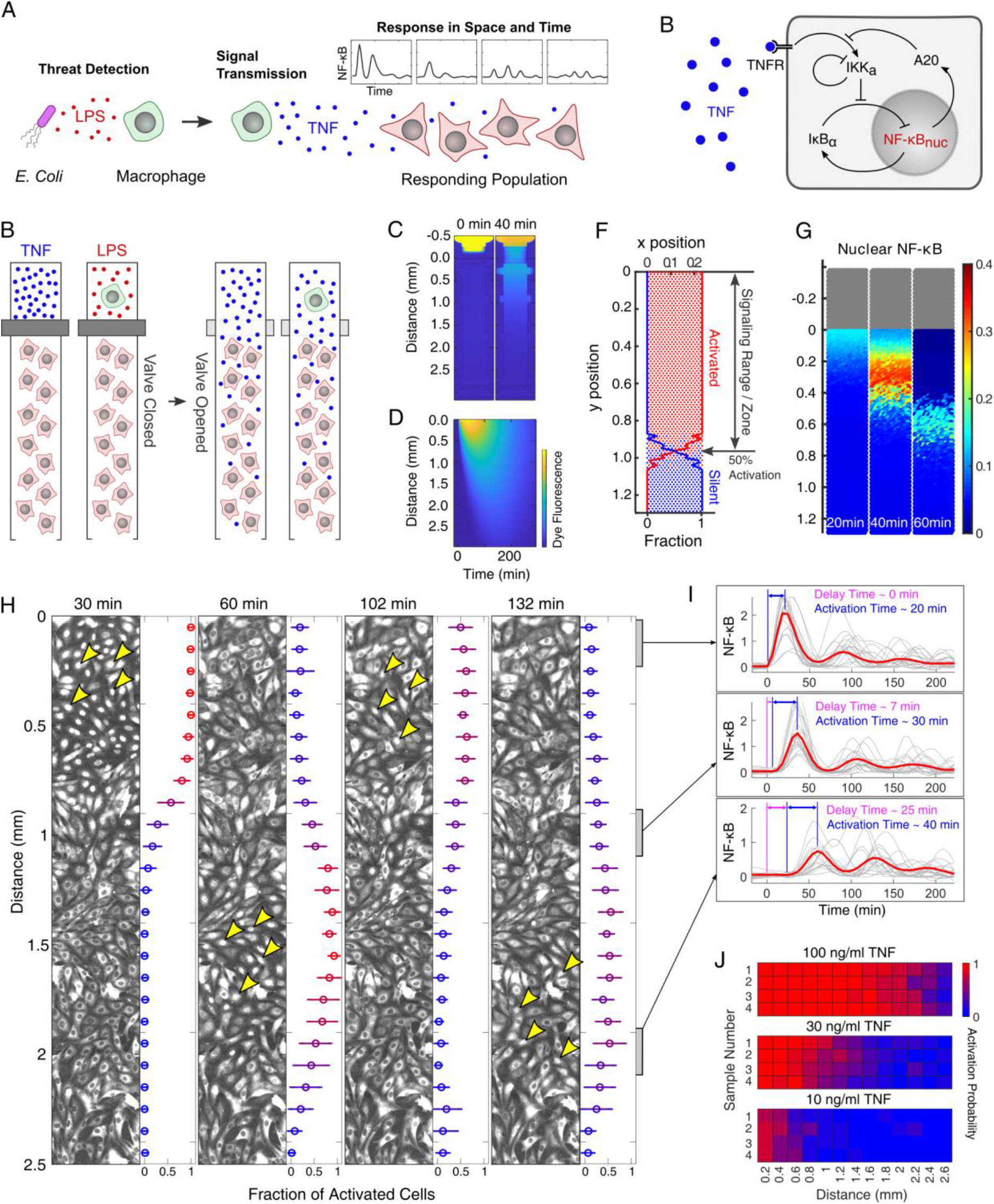
Cytokine secretion from a local source induces wave-like NF-κB propagation and a finite cell activation zone in the responding population. (A) Diagram illustrating a model of immune signaling following a local infectious signal. A macrophage detects a pathogen, and releases TNF to alert nearby cells. Receiving cells interpret this signal and regulate NF-κB dynamics. (B) Simplified NF-κB network used for mathematical modeling. (C) Microfluidic device designed to study the infection scenario represented in (A). Either TNF (blue dot) or a single macrophage was loaded in the source chamber. The macrophage was stimulated with LPS and washed with fresh medium. When the valve is opened, TNF or macrophage secretions diffuse into the receiving chamber and activate fibroblasts. (D) Diffusion of signals in the receiving chamber is tested with fluorescent dye. (E) Kinetics of dye fluorescence are plotted at various distances from the source chamber. (F) NF-κB activation at each location derived from simulation. Cell activation probability was calculated at various distance from source. Each point represents a single cell with the regulatory network in (C). (G) Simulation shows the release of TNF from source chamber elicits wave-like NF-κB propagation in the receiving cell population. (H) Microfluidic experiments exhibited wave propagation of NF-κB activation after release of TNF (100 ng/ml) from a source chamber. Fluorescently tagged NF-κB shuttles into the nucleus upon activation. Yellow arrows indicate examples of activated cells in each microscopy image. Graphs on the right of each image illustrate the fraction of activated cells at different distances. Circles and error bars indicate the mean and standard deviation in ten replicates. (I) NF-κB responses in near (0 – 0.4 mm), mid (0.8 – 1.2 mm), and far (1.8 – 2.2 mm) regions. Red lines show the mean of all cell responses in each region (n > 300), while thin lines represent 20 random responses. Purple bars indicate the delay time between the initiation of TNF diffusion and onset of local NF-κB activation, while blue bars indicate the activation time. (J) Activation probability at various distance calculated for three different doses of TNF (10, 30 and 100 ng/ml). Each dose had four replicates.

## Results

We computationally and experimentally considered a localized infection scenario that creates a one-way communication channel from a signal sending macrophage to signal receiving fibroblasts (Figure 1A). First, pathogen molecules (bacterial LPS) activate a macrophage in the infection site. The macrophage releases TNF that diffuses to neighboring cells and activate the NF-κB pathway. NF-κB regulates gene expression in a spatially and temporally varying manner in the responding population. We primarily focused on TNF signaling, as prior work showed that TNF is the main secreted factor from the RAW macrophages used in this study (Frank and Tay, 2015). This result was further confirmed by measuring the cytokine secretions from these cells (Figure S1). Upon stimulation by LPS, macrophages rapidly secreted TNF increasing by almost 100-fold, compared to less than 4-fold for all other measured cytokines. Furthermore, fibroblasts used in this study do not release signaling factors that activate NF-κB in other cells (Figure S5). This ensures that the communication scenario we created is indeed one-way, from the macrophage to the fibroblast population.

### Spatio-temporal modeling of NF-κB activity in a one-way communication scenario

To theoretically study spatial NF-κB signaling in our infection scenario, we built a simple model where TNF is released from a point source and allowed to diffuse freely through a long chamber filled with signal receiving cells (Figure 1C). The response of each cell was simulated according to an ordinary differential equation model of NF-κB dynamics in response to TNF (Figure 1B). To capture how variable activation affects signaling, each cell was assigned a random activation threshold (normal distribution, covariance (CV) = 0.2). This spatio-temporal mathematical model allowed us to probe how local TNF release produces a spatially and temporally resolved NF-κB response in a neighboring cell population (Figure 1F, G, S2 – 3).

Our modeling studies presented two interesting observations: the presence of a well-defined NF-κB activation zone in the responding population, and wave-like propagation of NF-κB activation. Despite single-cell activation variability, a well-defined NF-κB activation zone was created in the responding population, with a narrow boundary region between activated and non-responsive cell populations (Figure 1F). Within this zone, NF-κB activity spread in a wave-like manner, where a band of NF-κB activation propagated away from the TNF source (Figure 1G).

### Microfluidic experiments show a defined activation zone and wave-like NF-κB propagation in response to a cytokine source

To experimentally study spatiotemporal NF-κB response to local cytokine release, we developed a microfluidic device that can precisely control cytokine release from an isolated (source) chamber (Figure 1C) (Frank and Tay, 2015). To activate fibroblast cells, either soluble TNF or a single macrophage is first loaded in the source chamber. Then, a PDMS membrane valve is opened to allow the soluble TNF or macrophage secretions to diffuse into a receiving chamber filled with fibroblast cells (Figure 1D, Video SV1). The bottom end of the receiving chamber was continuously replenished (flowed) with fresh medium to simulate the role of capillaries in supplying fresh nutrients and depleting cytokine signals by providing a sink effect (Kumar et al., 2019). Our device thus simulates a localized infection and subsequent innate immune signaling scenario in tissue.

To track NF-κB response at the single cell level, we used 3T3 mouse embryonic fibroblasts expressing NF-κB fusion reporter (p65-dsRed). Fluorescence images reporting NF-κB nuclear localization were captured every 6 minutes, then analyzed with custom image analysis program to quantify NF-κB response in individual cell (Kellogg et al., 2014; Kudo et al., 2018; Lee et al., 2009).

Our live-cell imaging experiments supported the qualitative predictions from our mathematical model. When TNF was released from the source chamber, fibroblast population showed wave-like propagation of NF-κB activation in the receiving chamber (Figure 1H, Video SV2). Receiving cells were exposed to different cytokine kinetics depending on their distance to the signal source (Figure 1E). Cells at farther distance experienced slower increase in TNF concentration and lower maximal dose (Figure S4), which also delayed the onset of NF-κB activation (purple interval in Figure 1I) and extended the activation time (blue interval in Figure 1I) (Son et al., 2021; Tay et al., 2010). When these shifts are coupled with the oscillatory characteristics of the NF-κB network, it prompts a robust wave-like spread of NF-κB activation in the responding population.

Wave-like propagation could theoretically be due to signal secretion and amplification by responding cells (Danino et al., 2010; Hassinger et al., 1996). However, when we stimulated a subpopulation of fibroblasts with TNF, nearby fibroblasts did not show NF-κB activation, indicating that these cells do not secrete NF-κB activating signals (Figure S5). Thus, our results indicate that diffusion of TNF coupled with intrinsic features of the NF-κB network is sufficient to produce wave-like activation in the receiving cell population.

Our experiments also reproduced the sharply defined activation zone predicted by simulations (Figure 1J). For all tested source doses, the activation rate of receiving cells transitioned from > 75% to < 25% in less than 0.4 mm distance (< 15% of the chamber length). Furthermore, the signaling range (defined as a distance where 50% of cells are activated) varied minimally between culture replicates with the same source dose. These results indicate that the NF-κB signaling range is precisely defined, despite variable single cell activation inherent to the NF-κB network (Tay et al., 2010). These findings motivated us to study how the signaling range is affected by different cytokine secretion patterns from the source.

### Dose and duration of a cytokine signal independently control the range of signaling and number of NF-κB oscillations in receiving cells

Cytokine secretion from immune cells reflects the information about cell type, its current state, and the severity of the infectious challenge (Abasıyanık et al., 2020; Junkin et al., 2016; Rao et al., 2010). Different secretion patterns can potentially affect NF-κB signaling range and dynamics in receiving cells. In particular, previous studies demonstrated that a high number of NF-κB oscillations preferentially induce genes which determine cell fate and cause epigenetic remodeling through expression of late-term target genes (Cheng et al., 2021; Tay et al., 2010). Thus, we investigated how particular cytokine release parameters (e.g. dose and duration) affect the signaling range and NF-κB dynamics in nearby cells.

Returning to mathematical modeling, we adjusted the dose and duration of cytokine secretion from a point source and evaluated how secretion pattern affects the spatial NF-κB response (Figure 2A, B). Our simulation suggests that spatial features of NF-κB response to the same amount of cytokine depends primarily on the rate of release. Slow TNF secretion (Figure 2B, red) resulted in a short signaling range (distance from source) but a high number of NF-κB oscillations within the range. On the other hand, fast TNF secretion (Figure 2B, blue) showed long signaling range but low number of oscillations. When we only varied the dose of TNF secretion while keeping the duration constant, the signaling range linearly correlated with the logarithm of the source dose, but the number of oscillations was not affected (Figure 2C). On the other hand, the number of NF-κB oscillations correlated to the duration of TNF release, but was independent of the dose. Varying source dose did not affect the number of NF-κB oscillations, nor did varying duration affect the signaling range. Thus, our results indicate that dose and duration are two independent aspects of cytokine release that control different parameters of NF-κB response in signal receiving cells.

**Figure 2.**
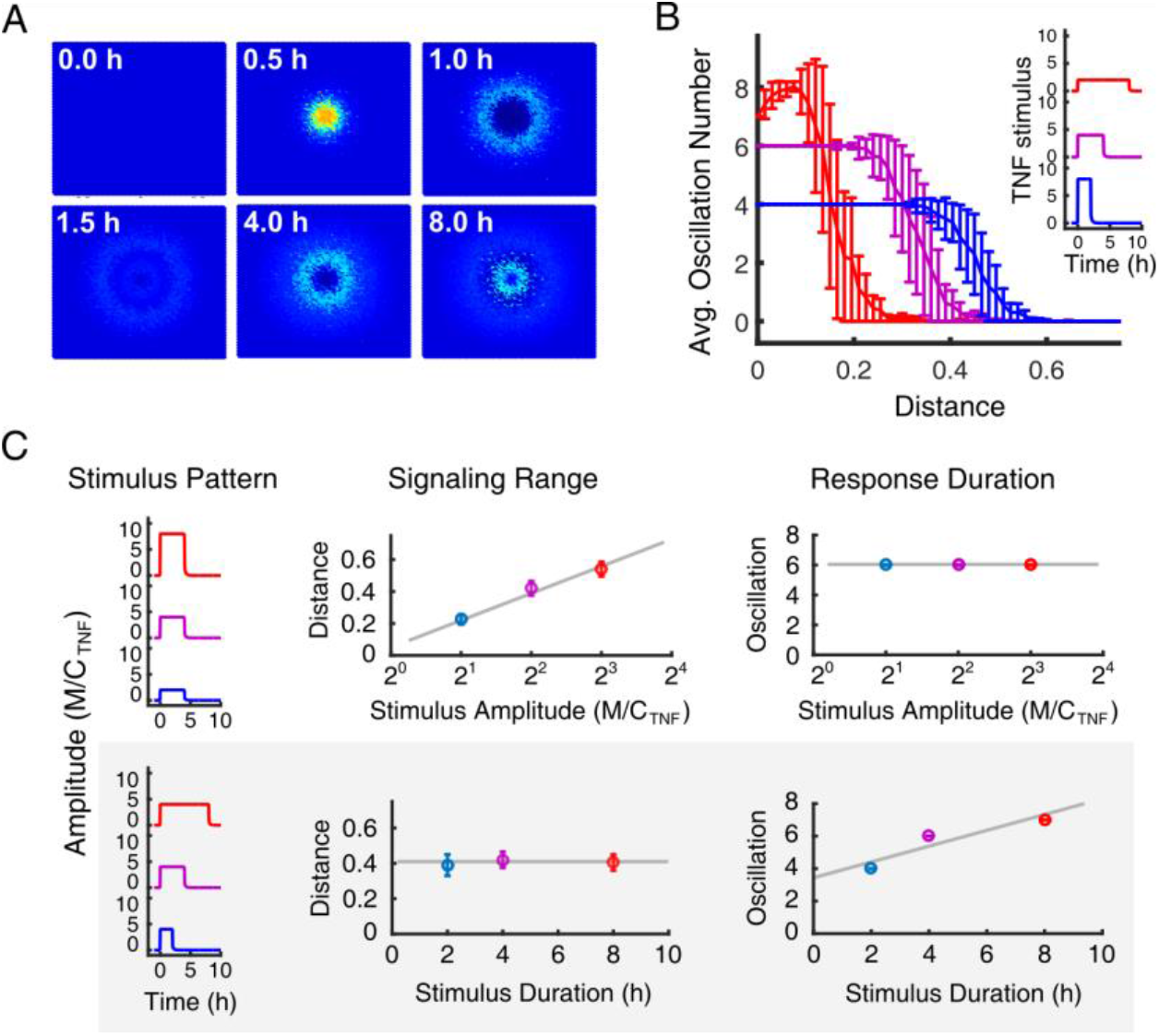
Modeling predicts that cytokine secretion dose and duration controls NF-κB response in distinctive and independent manner. (A) NF-κB activation in a 2-D tissue is simulated in response to TNF release at the center. (B) Spatiotemporal NF-κB responses to different secretion profiles with varied amplitude and duration are simulated. The total amount of cytokine or area-under-curve (AUC) in each profile is fixed for comparison. Blue indicates the population response for short and high release, while red indicates the response for long and low release. (C) From simulated results, the effect of secretion amplitude and duration on signaling range and number of oscillations is evaluated. Secretion profiles (stimulus pattern) and corresponding responses (signaling range and response duration) are grouped by color. Between top and bottom panel, the same color indicates stimuli with the same AUC.

We experimentally tested these predictions using our microfluidic system. When various doses of TNF (10, 30, or 100 ng/ml) were released from the source chamber, the signaling range increased proportionally to the logarithm of the source dose (Fig 3A, C). Similarly, when the TNF stimulus was released for different durations (15, 30, and 60 min), the number of NF-κB oscillations in the activation zone increased at longer stimulus durations (Figure 3B, C, S6). Regardless of the release duration, we observed a similar signaling range, with only ∼ 10 % increase in the range between 15 and 60 min. However, we observed a small increase in the number of oscillations depending on the source dose, despite our computational prediction that the two are independent (Figure 3C). This disagreement could be due to experimental depletion of TNF in the source chamber (stimulus patterns in top panel of Figure 3C), unlike the pulsatile stimulus used in our simulation. Regardless, we note that the number of oscillations was over twice as dependent on stimulus duration compared to stimulus dose. Overall, our simulation and experiments demonstrate that the signaling range strictly depends on the dose of stimulus, while number of oscillations depends significantly more on the duration of the stimulus.

**Figure 3.**
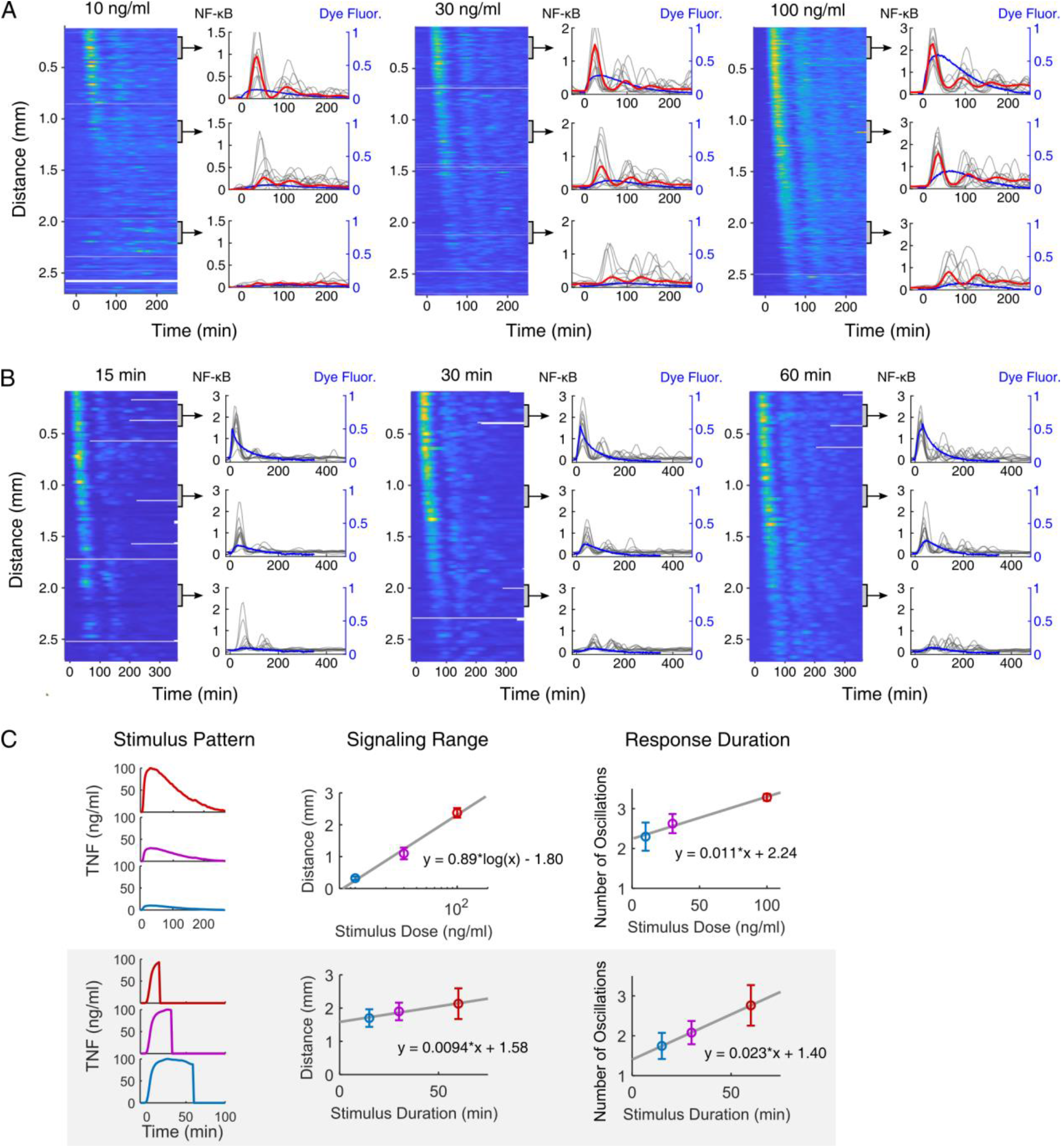
Dose and duration of cytokine secretion independently control the NF-κB signaling range and number of oscillations, respectively. (A, B) Heatmaps and traces depict the spatiotemporal NF-κB response to the TNF release from a source. Experiments were conducted with three different doses of TNF (10, 30, or 100 ng/ml) or by closing the separating valve 15, 30, or 60 min after releasing 100 ng/ml TNF. The plot on the right shows the NF-κB behavior in three different regions (top, mid, and bottom) in the cell chamber. Red lines show the mean behavior, while thin gray lines show ten random dynamics in each region. Blue lines show TNF dynamics predicted from the dye diffusion in Figure 1D. (C) Similar to Figure 2C, the effect of different stimulus patterns on the signaling range and number of oscillations is evaluated using the experimental data from (A) and (B). Data are displayed as mean ± SD from three replicates

### NF-κB independently decodes the dose, duration, and distance of a cytokine source

We investigated whether the NF-κB dynamics in signal receiving cells can independently decode the three key features of the signal source: the initial dose and duration of cytokine secretion, and its distance. First, to study how NF-κB dynamics change over distance, we organized the measured single cell responses based on their distance to the source chamber (Figure S7 – 8). As noted earlier, some differences, such as higher first peak or faster activation for closer cells, were due to the cytokine gradient formed by diffusion along the receiving chamber (Figure 1C). However, we also observed other parameters, such as the height and width of later peaks and the number of oscillations, significantly varied in a manner that cannot be attributed to the gradient alone (Figure 3A, S3, 9 – 11). In particular, cells located closer to the source had narrower and higher initial peaks followed by sharply decreased second peaks, while cells located farther had lower initial peaks but stronger second peaks.

To examine if these differences in the first and second peaks are sufficient to identify cell’s distance to the cytokine source, we quantified the peak heights and widths from each cell and clustered cells based on similarity in these parameters (Figure 4A, B, see S12 for 30 ng/ml TNF dose). When the cell positions in each cluster were plotted, we found that cells in each cluster were located at a similar distance from the TNF source (Figure 4C, S12). The single cell distances in each cluster were significantly different from the distances in the other clusters (Figure S13). Thus, the peak height and width within NF-κB dynamics contain enough information to define the receiving cell’s distance to the signaling source.

**Figure 4.**
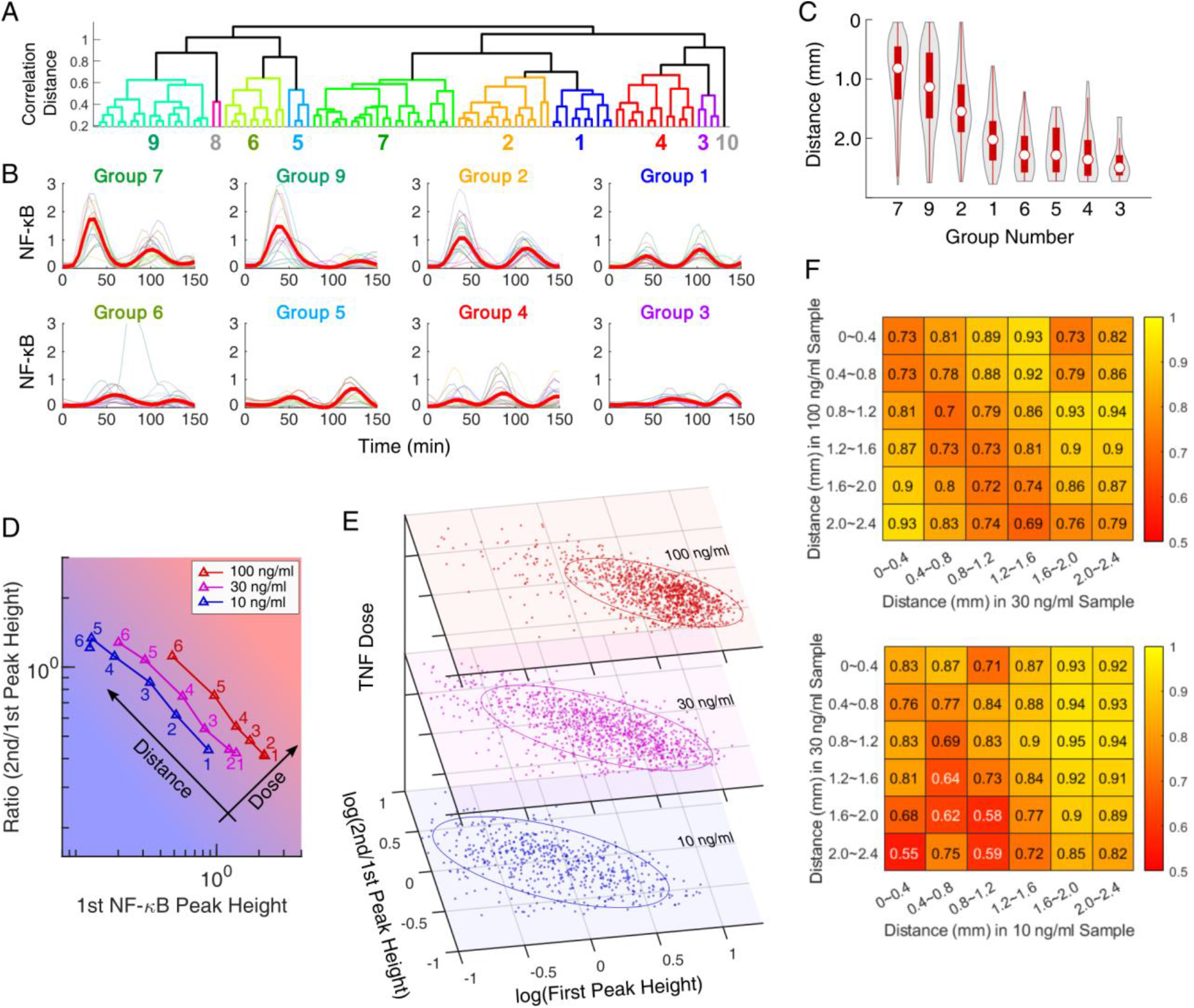
Dose, duration, and location of the cytokine source can be simultaneously determined from the NF-κB dynamics in the receiving cells. (A) Dendrogram shows the NF-κB dynamics in response to 100 ng/ml source dose divided into ten clusters. (B) Subplots show the NF-κB response behavior in eight major clusters. Thin lines show twenty random cell traces. Subplots were arranged based on the distance from the TNF source. (C) Violin-plots quantifying the spatial distribution of cells in each cluster. Red boxes indicate 25 and 75 percentiles, while circles represent the median distance. See Figure S13 for the statistical comparison. (D) Triangles show the mean first peak height and the mean ratio between first and second peak heights at six different distances from the cytokine source. Number on each point represents the relative distance to the source with one being closest. (E) Single cell data from different source doses are plotted in the same format as in (D). The ellipses indicate 20 % quantile isoline. (F) The single cell accuracy in distinguishing source dose and location is evaluated through cross-validation of the decision tree. The numbers in heatmaps indicate the distinguishability between the single cell responses in two locations from different source doses. Here, 1 indicates total distinguishability, while 0.5 indicates total indistinguishability.

We then examined if the source cytokine dose can be also identified from the NF-κB dynamics alone. Comparison of the dynamics at the same distance from different source doses shows that the height of the first peak increased with the source dose (Figure 3A). In contrast, the ratio between the first two peak heights specifically depended on the cell’s distance to the cytokine source. Irrespective of source dose, the ratio of the second peak height to first peak height increased with distance from the cytokine source (Figure S14). This result still held true when the area under the curve (AUC) of the peaks were compared instead of peak heights (Figure S14). Thus, a simple comparison of dynamic NF-κB features can elucidate two distinct attributes of the signaling source over distance.

Indeed, plotting the mean height of the first peak against ratio between the first two peaks revealed that cells from different distances line up along a single trajectory (Figure 4D, E, S15). More importantly, when single cell traces from different source doses were analyzed and plotted together, each dose formed distinct and parallel trajectories, implying that each distance and source dose can be differentiated from every other distance and source dose. Therefore, the cytokine dose and distance from the signaling source can be quantified independently, solely from the heights and ratio between the first two NF-κB peaks. In the previous section, we showed that the number of NF-κB oscillations in the receiving cells corresponds to the duration of the cytokine release. Therefore, our analyses collectively indicate that, if there are more than two oscillations present in the NF-κB dynamics, the dynamics contain sufficient information to simultaneously distinguish the dose, duration, and location of the cytokine source.

Although we find that the population mean of NF-κB dynamics carries multi-dimensional information about a cytokine source, the single cell responses vary significantly (Figure 4E, S7 – 11). To evaluate how accurately NF-κB dynamics estimate the location and dose of a cytokine stimulus in single cells, we classified single cell responses using binary decision trees. NF-κB localization traces from each source dose were aligned based on the onset timing of activation (Figure S16), and were divided into six groups based on their distance to the cytokine source. The accuracy of distinguishing between each group was quantified using 10-fold cross-validation (Figure 4F). For example, the number in the top left corner of the heatmap shows the distinguishability between the responses in first positions of two samples, where 1 indicates perfect distinguishability and 0.5 indicates total indistinguishability. Between the 30 and 100 ng/ml samples, groups were readily distinguished at the single cell level. All cells were classified with an accuracy of 0.69 to 0.94, even between the groups exposed to similar amount of cytokines, *e*.*g*., positions 4 and 5 with 100 ng/ml source dose vs. positions 3 and 4 with 30 ng/ml source dose (Figure S4). The distinction between groups still persisted when we compared cells from conditions with lower source doses, although the accuracy dropped to 0.55-0.94 due to higher relative noise at lower doses. Overall, despite noise in the NF-κB network, our analysis shows that individual cells can process the local cytokine dynamics and decode information about cytokine secretion dose and its relative location.

### Macrophage secretion variability determines NF-κB signaling range and duration

TNF secretion by macrophages varies significantly between individual cells even under identical pathogen stimuli (Baer et al., 1998; Junkin et al., 2016), but the consequence of cytokine variability on NF-κB signaling is not understood. To predict the effect of variable cytokine secretion on NF-κB signaling range, we used previously measured TNF secretion profiles from single macrophages as input to our stochastic model simulations (Figure 5A) (Junkin et al., 2016). We found little variation in the signaling range (the physical extent of NF-κB activity in the responding cell population) when the same secretion profile was tested multiple times, despite variability in the NF-κB response of receiving cells (Fig 5A top). However, using different secretion profiles from macrophages resulted in highly variable signaling range (Figure 5A bottom). Thus, the difference in cytokine secretion dynamics is readily translated into variation in the NF-κB activation range.

**Figure 5.**
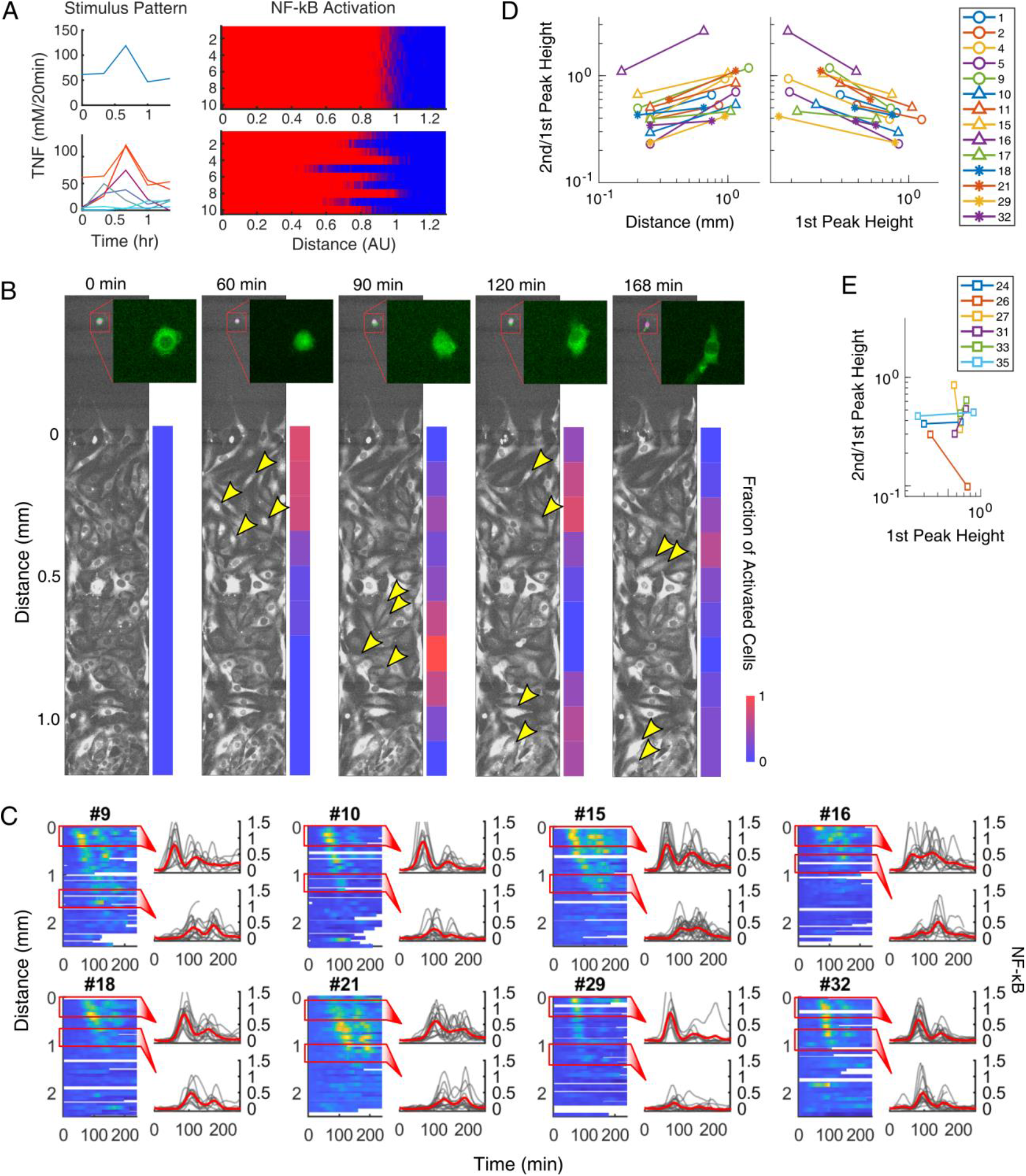
Macrophage cytokine secretion dictates the signaling range and duration. (A) Using measured TNF secretion profiles from macrophage, the effect of stochastic NF-κB response on signaling range is simulated. Top right shows the results from ten different trials using an identical secretion profile (top left). Bottom right shows the results from each secretion profile shown on the left. (B) Time-lapse fluorescence images show our co-culture experiments, which imitate the tissue infection. A macrophage harboring p65-gfp was stimulated with 10 ng/ml LPS (green cell in the inset), which released TNF to activate NF-κB in the receiving population (bottom chamber). Yellow arrows indicate examples of activated cells in each microscopy image. The color bar on the right shows the activation rate at various distance. (C) Heatmaps and single cell traces show ten examples of the spatiotemporal NF-κB response induced by single macrophages. See Figure S17 for all samples. (D, E) Similar to Figure 4D, the mean ratio between first and second peaks is plotted against the mean first peak height for near and far regions. Additionally, the relationship between distance and peak ratio is also compared to validate distance dependence of the ratio. Numbers in the legend correspond to the numbers in Figure 5C and S17. (D) shows samples with similar slopes, while (E) shows outliers.

To validate this prediction, we performed macrophage-fibroblast co-culture experiments which modeled inflammatory signaling from an activated macrophage to neighboring fibroblasts. A single RAW264.7 macrophage was loaded into each source chamber in our co-culture device, stimulated with 50 ng/ml LPS for 10 minutes, and washed with fresh medium to remove LPS. The separator valve was then opened to allow LPS-induced macrophage secretions to diffuse to fibroblast population (Figure 1B). This stimulation method induced strong NF-κB activation in most macrophages, although the duration of NF-κB activation varied widely (1 − 4 h) in each macrophage (Figure 5B).

The co-culture experiments also showed wave-like propagation of NF-κB activation through the receiving cells (Figure 5B, Video V3), similar to our observations from TNF diffusion experiments (Figure 1G). However, unlike TNF diffusion experiments, we observed a wide range of signaling distances and durations when activated macrophages were used as signal source (Figure 5C, S17). Our experiment with thirty-five samples, each with a single macrophage in the source chamber and one to two hundred fibroblasts in the receiving chamber, showed that the NF-κB signaling range varied from almost 0 to 2 millimeters. Likewise, the number of NF-κB oscillations in the receiving cells varied from 0 to more than 3, depending on each macrophage and location of the receiving cell (Figure 5C, S17). Compared to the minimal variation we found from earlier TNF diffusion experiments, different secretion patterns by macrophages had a significant influence on the signaling range and number of NF-κB oscillations in the surrounding cells. Thus, our results suggest that the local macrophage secretion patterns tune and directly control the local inflammatory response during infection.

We also highlight that the NF-κB pathway in fibroblasts was able to decode the cytokine dose and distance to the macrophage despite the variability in the secretion patterns. When the ratio between the first and second NF-κB peak were plotted against distance or against first peak height, roughly 70% of trajectories had similar slopes (slope CV < 0.5) (Figure 5D). While a minority of experiments displayed aberrant NF-κB dynamics (Figure 5E), the majority of samples behaved similarly. Hence, despite variability in single macrophages, NF-κB dynamics retain the ability to encode the receiving cell’s relative location and the dose of cytokine secretion from macrophage.

### Cytokine dynamics determines spatial gene expression patterns in the responding population

Although we demonstrated that the NF-κB dynamics in receiving cells can encode the distance and dose of the secreting source (Figure 4), how these differences in dynamics affect the transcription of response genes is unclear. Previous studies suggest that mRNA degradation rate is a key parameter in understanding different kinetics of NF-κB target genes (Dorrington and Fraser, 2019; Sen et al., 2020; Son et al., 2021; Tay et al., 2010). Genes involved in NF-κB feedback, like the IκB family and A20, are rapidly expressed upon TNF stimulus, but also rapidly degraded (early genes). On the other hand, inflammatory response genes, *e*.*g*., those related to apoptosis or chemokine signaling, show slower activation and prolonged expression afterwards (late genes). We hypothesized that each NF-κB target gene would show a distinct expression pattern over the activation zone, depending on its dynamic characteristics and the responding cell’s distance to the cytokine source.

To estimate how local gene expression profiles change at different distances from the signaling source, we added a simple gene expression component to our mathematical model and varied the degradation rate of mRNA (Figure 6A, Method). Early genes closely followed NF-κB localization dynamics, rapidly diminishing upon NF-κB inactivation. As a result, expression of early genes is restricted to the vicinity of the source, where NF-κB oscillations persisted for longer time. On the other hand, late genes slowly accumulated during NF-κB activation and remained high even after NF-κB inactivation. When the duration of stimulus was long enough, the difference in gene expression characteristics resulted in a spatial pattern, where early genes were upregulated in vicinity of the cytokine source while late genes rose at longer distances.

**Figure 6.**
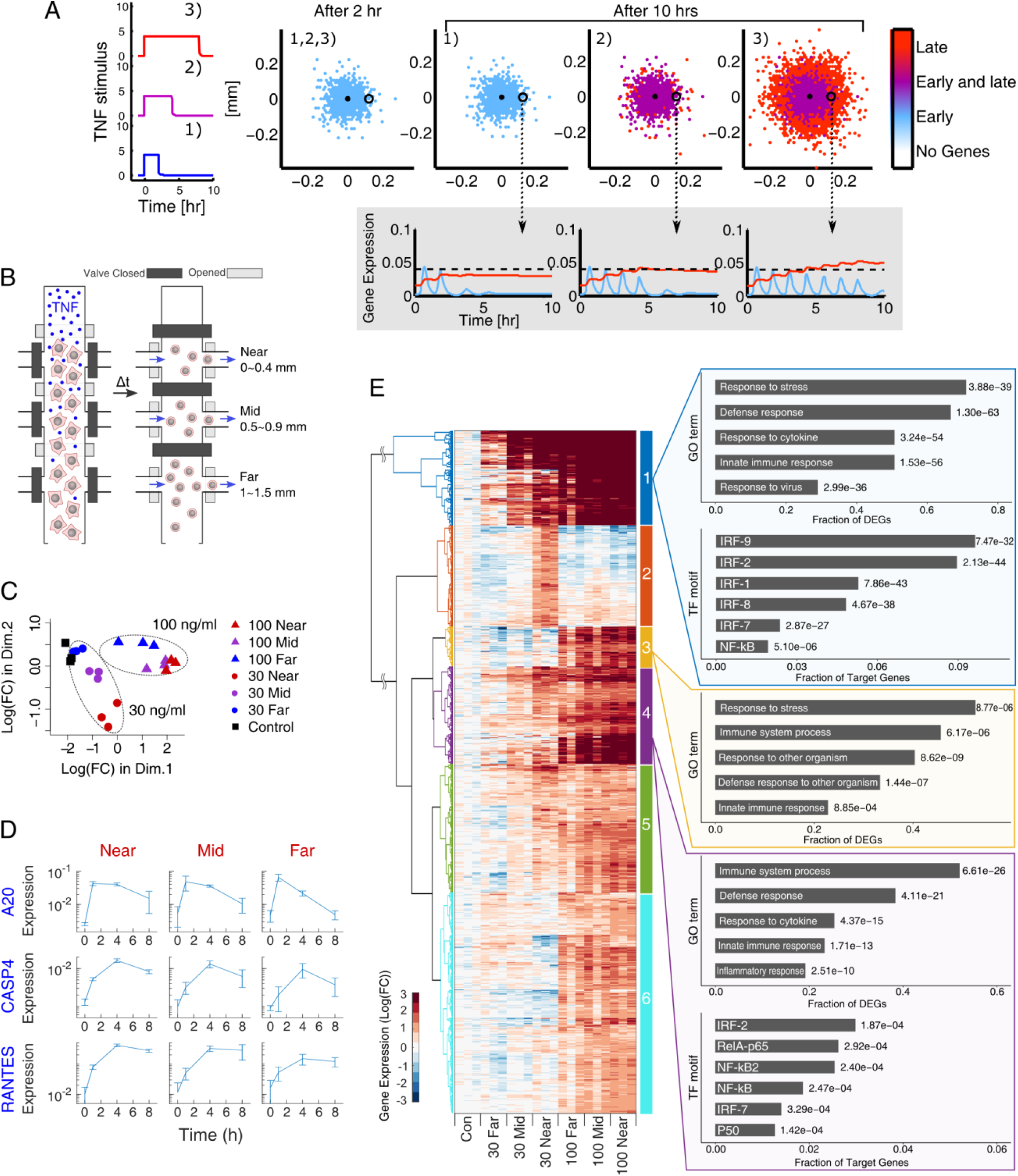
Spatially resolved gene expression analyses highlight distinct pro-inflammatory gene clusters depending on distance and source dose. (A) Downstream expressions from three different cytokine secretion durations (2, 4, and 8 h) are simulated with our 2-D model. Expression at each grid point is calculated using the corresponding NF-κB dynamics as an input to a gene expression function. Blue indicates genes with rapid degradation (early), while red represents genes with slow degradation (late). (B) Diagram illustrates the schematics of spatially-resolved downstream measurement using a custom microfluidic chip. (C) Cells from three different regions are collected 4 h after releasing TNF from source chamber (30 or 100 ng/ml). After sequencing, the pairwise distance between samples are compared in MDS plot for 1,094 significantly upregulated genes. (D) Using RT-qPCR, the kinetics of A20 (early), CASP4, and RANTES (late) genes are measured for different locations in 100 ng/ml sample. (E) Upregulated genes from sequencing data were clustered using correlation method. The functional characteristics of genes in each cluster were analyzed through enrichment of GO terms and TF (transcription factor) motifs. The numbers to the right of the bar indicate the *p*-value.

We tested this prediction by measuring gene expression from three different regions in the receiving chamber in TNF diffusion experiments (Figure 6B). Using a custom microfluidic device, cells were retrieved from all three regions (and at multiple time points: 0, 2, 4 and 8 h) following exposure to the spatial gradient formed by a 100 ng/ml source dose. RT-qPCR was used to measure the kinetics of key early and late NF-κB response genes in the responding population from each three locations (Figure 6D) (Tay et al. 2010). Early genes (*A20*) were rapidly induced and degraded, while late genes (*Casp4/RANTES*) rose and fell more slowly. These trends led to spatial patterning of gene levels at later time intervals. At 8 h, *A20* remained highly expressed near the source but sharply decreased at farther distances. In contrast, *Casp4* and *RANTES* were highly expressed in all three regions (near, mid, and far). Combined, our experiments showed spatial patterning of early and late response genes in the responding population. In summary, we found that distinct spatiotemporal gene expression profiles can arise from a combination of gene-specific degradation rates, single-cell NF-κB dynamics, and cytokine diffusion from a point source.

### Spatially resolved RNA sequencing shows distinct pro-inflammatory gene clusters at different distances from the cytokine source

RNA sequencing was used to more closely investigate the spatial variability in the gene expression patterns. Cells were stimulated with two different source doses (30 and 100 ng/ml TNF), and retrieved after 4 hours, where expression levels were close to the maximum for all three genes in our time-course measurement (Figure 6D). Among all genes evaluated, 1,094 genes were significantly upregulated compared to the control (no exposure to TNF), and used for further analysis.

First, to see if the expression of response genes is differentiated by the receiving cell’s location and source dose, we calculated the pairwise distances between the samples and visualized them with multidimensional scaling (MDS) (Figure 6C) (Anders et al., 2013; Robinson et al., 2010). In the scatter plot, we noticed that samples grouped based on the source dose. In particular, the mid and far 100 ng/ml samples were clearly separated from the near 30 ng/ml samples, even though the maximum cytokine concentrations they experienced during the experiment were similar (Figure S4). This indicates that genes are differentially regulated based on both location and source dose.

We then performed hierarchical clustering to group genes based on their dependence on the source dose and distance, and processed each group through gene ontology (GO) analysis to analyze the functional characteristics of each gene cluster (Figure 6E, Table ST1 – 12) (Ashburner et al., 2000; Raudvere et al., 2019). Clustering resulted in six groups with different patterns of upregulation. In particular, cluster 1 (blue) genes were strongly induced at most distances in both source doses, cluster 2 (red) genes were expressed specifically in the top position of 30 ng/ml, cluster 3 (yellow) genes were strongly expressed in the 100 ng/ml but not the 30 ng/ml source dose, and cluster 4 (purple) included genes that showed strong dependence on position and dose. The GO term analysis shows that cluster 1, 3, and 4 are enriched with genes related to the immune or stress response, which is expected from TNF stimulus. However, genes in cluster 1 were mostly regulated by the interferon regulatory factor (IRF) family of transcription factors, while genes in cluster 4 were largely regulated by the NF-κB transcription factor family. Since genes in cluster 1 were strongly upregulated in most positions, this result suggests that IRF-dependent genes are highly sensitive to TNF stimulus and that IRF can drive long-range effects during inflammatory signaling. In contrast, NF-κB-dependent genes were mostly upregulated close to the source, possibly due to the higher threshold for activation by TNF (Figure 7). Thus, the difference between IRF and NF-κB target gene expressions would increase at farther distances, producing spatial variability in inflammatory gene expression patterns. Surprisingly, some immune genes are almost independent of the distance (cluster 3). Even though the maximum cytokine concentration was higher in the near position of 30 ng/ml than the far position of 100 ng/ml (Figure S7), immune genes in cluster 3 were strongly expressed in all positions of 100 ng/ml while hardly expressed in any of 30 ng/ml sample. These data show that dose-dependent but distance-independent signaling is possible within the signaling zone (Figure 7). The other clusters (cluster 2, 5 and 6) showed upregulated genes related to cell growth and metabolism as well as stress response (Table ST4, 10, 12). Overall, our sequencing analysis demonstrates different roles for two transcription families in inflammatory signaling, and classifies immune genes which are regulated based on the distance to the source and/or the source doses.

**Figure 7.**
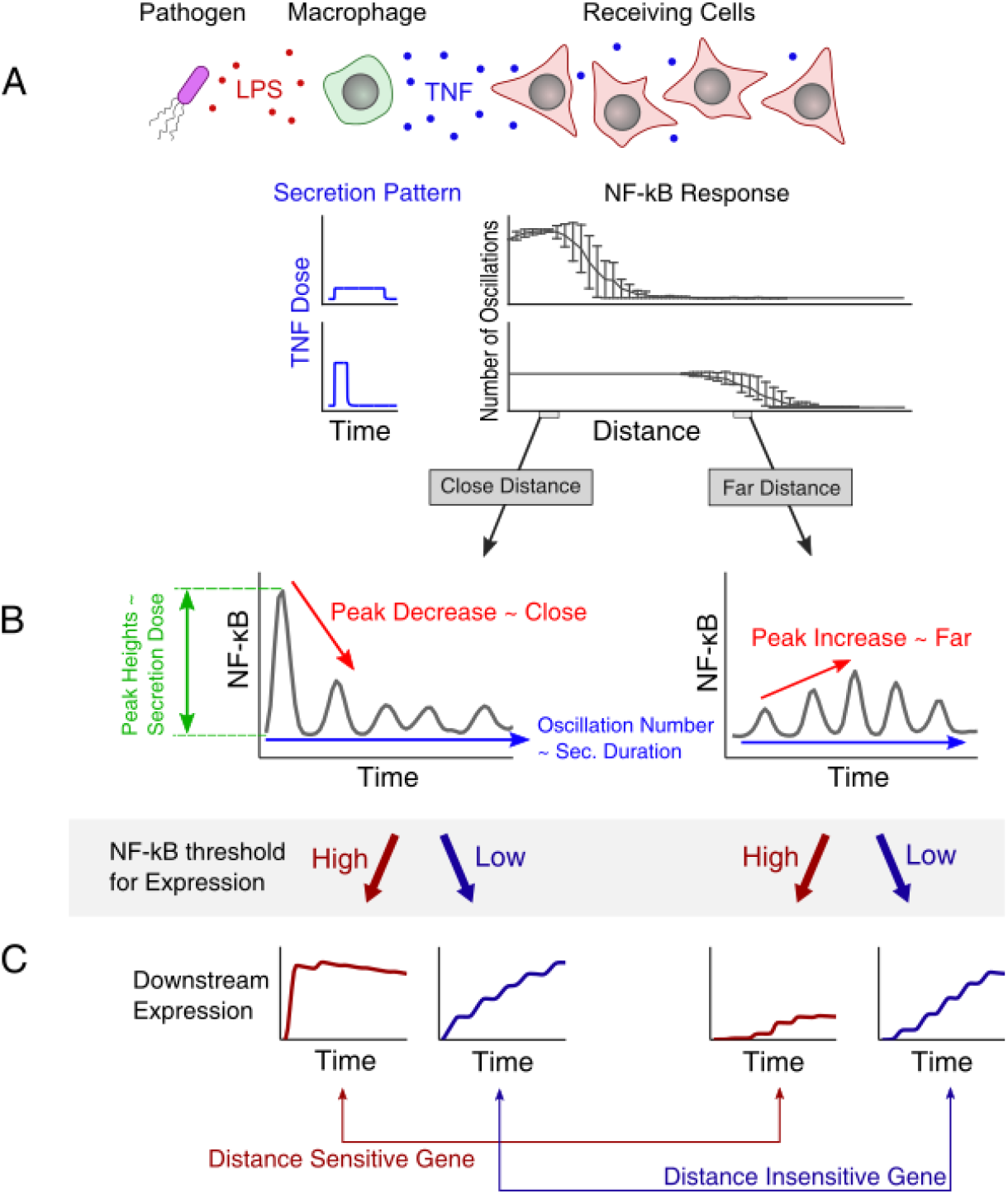
Cytokine secretion pattern determines NF-κB activation range and duration, and responding cells regulate distance-specific inflammatory responses. (A) The dose of cytokine secretion by a macrophage controls signaling range, while the duration of secretion controls the number of oscillations in the responding cells. (B) Gray lines represent single cell NF-κB dynamics at close and far distances, while colored arrows highlight the different features of NF-κB dynamics corresponding to different attributes of cytokine secretion by the macrophage. (C) Using the NF-κB dynamics in (B) as input, the kinetics of two downstream genes (red and blue) with different NF-κB thresholds are simulated.

## Discussion

Inflammatory signals in response to infection are often initiated from a localized point in the cellular microenvironment. In this study, we used the NF-κB network to explore how localized cytokine secretion affects spatiotemporal aspects of pro-inflammatory signaling at the single cell and population levels (Figure 7A). Through mathematical modeling, microfluidic co-culture experiments and spatially resolved gene expression analysis, we showed that localized cytokine secretion from a macrophage creates wave-like propagation of NF-κB activation in responding population. The range of signal propagation is sharply defined and strictly correlated to the dose of cytokine release, irrespective of the release duration. On the contrary, the number of NF-κB oscillations in the signaling zone depends on the release duration. Hence, the signaling range and duration of NF-κB response in a cell population are two independent properties that can be attributed to dose and duration of cytokine secretion. Stochastic regulation of the NF-κB network minimally affected these properties of spatial response. Hence, each macrophage can tune its cytokine secretion pattern to control the range and duration of the inflammatory response in the neighboring cells. Surprisingly, NF-κB dynamics can independently encode the dose, duration, and location of a cytokine source (such as a macrophage), and gives rise to distinct gene expression patterns depending on the cytokine secretion dose and distance. Overall, our study highlights the ability of NF-κB signaling network to gain detailed information about a signal source by interpreting the kinetics of diffusing cytokines, and coordinate distance and source specific responses in a cell population (Figure 6, 7B – C).

RNA sequencing analysis from different locations in the responding cell population revealed a group of immune genes that showed strong correlation to the cytokine dose rather than its distance. Since cytokine secretion dose is correlated with the severity of infection (i.e. the pathogen load (Iwasaki et al., 2010)), this proposes that receiving cells can interpret the severity of infection from the diffusing signal even at a distance. Such a capability would allow local cells to coordinate their inflammatory response based on the distance and severity of the infection to accurately interpret and overcome infectious challenges.

Additionally, we found IRF-family genes are more sensitive to TNF than NF-κB-family genes, resulting in significantly longer signaling range for IRF target genes. IRF is closely associated with antiviral innate immunity, and previous studies have shown its crosstalk with the NF-κB pathway (Czerkies et al., 2018; Rubio et al., 2013). However, IRFs can also control distinct inflammatory pathways (Kaisho and Tanaka, 2008), and recent studies have shown that differential regulation between NF-κB and IRF promotes different inflammatory responses (Bender et al., 2020). Since our analysis showed that the differential regulation between IRF and NF-κB pathway is enhanced at farther distance, these results indicate that different activation thresholds for two distinct transcription factors can be used to fine-tune distance specific inflammatory responses.

## Acknowledgements

This work is supported by NIH Grants R01GM128042 and R01GM127527, and Army Research Office grant 73231-EL (S.T.). A.T. received funding from the Danish National Research Foundation (grant number: DNRF116).

## Author Contributions

M.S. and T.F. designed and performed the microfluidic experiments with help from M.J., A.G.W., and S.S.K. M.S., T.F., T.H.H., and A.T. analyzed the data. T.H.H and A.T. built the mathematical models and performed the simulations. M.S. and A.G.W. designed and performed the downstream measurements and data analysis. S.T. supervised the work. All authors contributed to writing of the manuscript.

## METHODS

### RESOURCE AVAILABILITY

#### Data and code availability

- RNA sequencing data have been deposited at NCBI Gene Expression Omnibus (GEO) database and are publicly available as of the date of publication.
- Single cell NF-κB dynamics for all experiments and codes for heatmaps and single cell dynamics have been deposited in the github (https://github.com/tay-lab/Spatial_NF-kB_Signaling) and are publicly available as of the date of publication.
- The code for the MDS plot and clustering of RNA sequencing data have been deposited in the same github address as above.
- Any additional information required to reanalyze the data reported in this paper is available from the lead contact upon request.

#### Lead contact

Further information and requests for resources and reagents should be directed to and will be fulfilled by the lead contact, Savaş Tay.

## METHOD DETAILS

### Cell culture

Mouse fibroblast (p65-/- NIH 3T3) harboring H2B-GFP (nucleus marker) and p65-DsRed (NF-κB marker) was cultivated using Fluorobrite DMEM (Gibco, A1896701) supplemented with 10% FBS (Omega Scientific, FB11), 1% GlutaMax (Gibco), and 1× penicillin-streptomycin (Gibco) (Lee et al., 2009; Tay et al., 2010). For passaging, cells were harvested by adding Trypsin-EDTA (Gibco), washed, then re-suspended in fresh medium (1:10) before reaching 100% confluency in regular flask. For macrophage (p65-/- RAW 264.7) harboring p65-GFP reporter, we used the same DMEM medium supplemented with 10% certified FBS (Gibco, A3160401), 1% GlutaMax (Gibco), and HEPES (Gibco, 15630106). Before reaching 100% confluency, cells were scraped and re-suspended in fresh medium (1:10).

### Microfluidic device fabrication

The patterns for our diffusion microfluidic chip were designed with AutoCAD (Autodesk, Inc.) and KLayout. To fabricate the relief mold with the design, we followed the standard soft lithography procedure. For the valve regions on the flow layer, we used positive photoresist (MicroChemicals, AZ40XT) with 20 μm thickness. For the other part of flow layer or control layer with valves, we used negative photoresist (MicroChemicals, SU-8 3025) with 30 or 25 μm thickness respectively. All photoresists were exposed to 375 nm laser using the maskless aligner (Heidelberg MLA150). After developing, all molds were hard-baked, which generated rounded cross-section for the positive photoresist features. For more details about the mold-making protocol, see Gómez-Sjöberg et al (Gómez-Sjöberg et al., 2007).

For PDMS casting, we first treated molds with chlorotrimethylsilane (Sigma-Aldrich, 92360) to prevent the bonding between the mold and PDMS. Polymer and catalyst for PDMS (Momentive, RTV-615) were mixed in 10:1 ratio. To create a slab of PDMS grooved with control valve pattern, the PDMS mixture was poured over the mold for control layer, then degassed in vacuum desiccator. For the control layer, 5∼10 g of mixture was poured on the mold, and spun at 2300 rpm for 1 min to create thin (∼50 μm) layer of PDMS. Both mixtures were cured overnight at 80 °C. Next day, the cured PDMS slab and thin layer were treated with oxygen plasma (Harrick, PDC-001) for 30 sec, then were aligned using a custom stage with upright microscope. For the robust bonding between the two layers, the aligned PDMS bodies were baked overnight again, then the holes for the control lines and fluid inlets were punched. The combined and punched chip was bonded to a clean glass slide through oxygen plasma treatment. More details about the fabrication process is described in our previous publication (Kellogg et al., 2014).

### Microfluidic experiment

Using Tygon™ tubing (ND-100-80) and metal pins, each valve in the control layer was connected to a pneumatic solenoid valve (Festo, 197334). The tubes were prefilled with water to prevent diffusion of gas to the fluid channels. The solenoid valve was controlled electronically, whose exact timing for opening and closing was controlled by custom developed Matlab interface. Various medium containing cells or ligand was aliquoted in a pressure relief vial and delivered to the microfluidic chambers through PEEK tubing (VICI®, TPK.505). The stage of microscope (Nikon Eclipse Ti2) was surrounded by a custom-made box to maintain the temperature at 37 °C (LifeImagingSecrvice, Basel), and the microfluidic device was enclosed in a stage-top-incubator, which maintained the chip environment with 5% CO_2_ and above 98 % humidity. More details about the setup and control over the chip were described in our previous publication (Kellogg et al., 2014).

For the fibroblast loading, the surface in microfluidic chambers was treated with 1:4 fibronectin (1 mg/mL) in PBS solution overnight, washed with fresh medium, then fibroblasts were loaded in the diffusion chamber at about 80 % confluency. After seeding, we waited at least 5 h with one or two feedings in the middle to allow cells to attach and spread uniformly in the diffusion chamber. In the meantime, cells stuck or settled in the other channels were removed by flushing with trypsin. For the macrophage loading in the source chamber, cells were diluted in the fresh medium, and the channel was opened and closed repeatedly until one cell was seated in the source chamber. All microfluidic chips were controlled by a custom graphic user interface (Matlab), whose underlying software architecture and hardware-software interface were modified from Gomez-Sjörberg et al (Gómez-Sjöberg et al., 2007). The software could also run a pre-written code which automatically feeds, sinks, and purges the microfluidic chambers at designated time.

During the diffusion experiment, the bottom channel of the diffusion chamber was continuously washed by flowing fresh medium through on-chip peristaltic pumping. To ensure consistent cytokine sinking without disturbing the medium inside the diffusion chamber, the peristaltic pumping was performed at very low frequency (0.33 Hz or 20 cycles/min) and the bypass was washed every 1∼2 min. This process also allowed the medium exchange by diffusion from the sink to the diffusion chamber and enabled cells to continuously grow without additional feeding. For the diffusion experiment with cytokine, either 10, 30, or 100 ng/ml TNF (R&D Systems, 410-MT) was loaded in the source chamber then the separating valve was opened, while the bottom sink channel was continuously pumped with fresh medium. For the diffusion experiments with different duration of TNF release (Figure 3B), 100 ng/ml TNF was loaded in the source chamber, the separating valve was opened, then closed after 15, 30, or 60 min. For the co-culture experiments with RAW macrophage cell, the macrophage was stimulated with 10 ng/ml of LPS (InvivoGen, tlrl-3pelps) for 10 min, washed with fresh medium, and the separating valve was opened.

### Image acquisition

Fluorescence time-lapse images were acquired with CMOS camera (Hamamatsu, ORCA-Flash4.0 V2) using LED as excitation light source (Lumencor, Spectra X). For the p65-DsRed imaging, cells were imaged with 555 nm excitation with 0.3 sec exposure time and 100 % LED intensity, while the nucleus marker (H2B-GFP) was captured with 470 nm wavelength and 40 msec exposure time at 100 % intensity. All images were acquired with a Nikon S plan Fluor ELWD 20x objective for both fluorescence channel. Images were acquired every 6 min using automated acquisition control by NIS element software (Nikon). Photo bleaching was not observed even after 100 sets of fluorescence images collected in this setup (equivalent to 10 h duration of the experiment). Images from each experiment were corrected by subtracting dark frame images and dividing them by the flat field images.

### Image and data analysis

With the magnification we used (20x), each diffusion chamber needed four frames or positions to capture fluorescence images of entire chamber. For each chamber and time point, these images were stitched using custom Matlab software which utilized ‘detectSURFFeatures’ and ‘estimateGeometricTransform’ functions to create the transform matrix. Then the stitched images were processed through a custom image analysis software (Matlab) to extract the NF-κB response behavior for each cell in the chamber. Briefly, the software first identified the location and boundary of nucleus through H2B-GFP fluorescence images. Integrating the positions and sizes of nucleus across a sequence of images, the trajectory of each nucleus was evaluated. To sort out the dead or dividing cells, the program also monitors change in the shape or contrast of nucleus, and deletes the trajectories that showed any abnormal change in the middle. The selected trajectory and boundary of the nucleus were applied to the corresponding p65-DsRed images to evaluate the NF-κB nucleus translocation, using the similar method as reported in the previous study (Kudo et al., 2018). The background fluorescence was estimated based on the mean fluorescence in a few small areas without cell, which was subtracted from the corresponding p65 images. The resulting NF-κB dynamics were smoothed using “lowess” method to reduce the noise from cell migration or collisions between cells. The NF-κB level measured prior to the start of the experiment (opening of the separating valve) was considered as basal level for all data in this study. For the analysis that required number of activated cells in each sub region, two standard deviations above the mean basal level was used as threshold.

### Downstream expression measurement

Cell retrieval process largely followed previously described protocols (Mehling et al., 2015). To facilitate the retrieval of cells, the outlet of PDMS chip was cut with razor, so that the purge medium carrying retrieved cells flowed directly onto the glass slide. This allowed cells to settle on the open space in the form of a droplet, which could be retrieved with simple pipetting. After stimulating cells with indicated time, cell chamber was washed and incubated with TryLE express (Gibco) to detach cells from the glass surface for 1 – 2 min. Then the separating valves were actuated to divide the cell chamber into top, middle, and bottom sections and isolate cells in each position. Each position was then purged with PBS to send detached cells to the outlet. Cells in a ∼2 µL droplet was removed and deposited in 10 µL ice cold lysis buffer containing 0.1% Triton-X-100 and RNAse inhibitor (Takara). Lysate was stored at -80°C until further processing.

For qPCR, gene-specific reverse transcription and pre-amplification were performed using established protocols using a CellDirect One-Step RT qPCR kit (Thermo Fisher). Custom primer/probe sets were used to measure the expression of Tnfaip3 (A20), Casp4, and Ccl5 (RANTES), whose sequences are listed in our previous publication (Son et al., 2021). Ct values were calculated using software defaults, and normalized to GAPDH to produce ΔCT. The fold-change or relative expression level was calculated by comparing ΔCT in stimulated samples to unstimulated cells, or simply using following equation, 2^(ΔCT_unstimulated_ – ΔCT_stimulated_).

For RNA sequencing, samples were processed through the previously established SMART-Seq2 pipeline with minor variations (Picelli et al., 2014). The first strand synthesis and template switching were performed using SuperScript II (Thermo Fisher) and a modified template-switching oligo (5′-AAGCAGTGGTATCAACGCAGAGTGAATrGrGrG -3′), which were followed by 10 cycles of pre-amplification using KAPA HiFi (Roche). The amplified samples were tagmented and indexed (library prep) using Nextera XT reagents (Illumina). The prepped samples were sequenced at the University of Chicago Genomics Facility using Illumina HiSEQ4000 system with a read length of 50 bp. Adapter trimming and read mapping were done using STAR with default parameters. Transcript abundance was quantified using featureCounts. Raw counts were normalized and differentially expressed genes were identified through corresponding R packages, edgeR and limma. Differential genes between samples were identified using cutoffs FDR < 0.01 and log-fold change > 1. Gene Ontology (GO) terms and transcription factor motifs were identified using the web tool g:Profiler.

## MATHEMATICAL MODELING AND STOCHASTIC SIMULATIONS

### System of equations for NF-κB network

To capture general behavior and dynamics of NF-κB translocation and associated components, we modified the previously published models, and fitted the parameters to the experimental data (Mengel et al., 2012). The core of our model was derived from the minimal model where they reduced the process of transport of NF-κB in and out of the nucleus from a seven-variable to a three-variable system (Krishna et al., 2006). From the minimal model, we applied following changes to qualitatively describe the observed behavior at both single cell and tissue level (Eq. M1 – 10).

- The activation term of IKK was changed from a linear term in TNF, to a non-linear Hill-term. The Hill-term contains an explicit activation threshold, C_TNF_, which is not included in models with linear activation. We normalized the TNF concentration with respect to the threshold, C_TNF_.
- To increase the accuracy of simulation, we added the upstream negative feedback (A20) induced by nuclear NF-κB and the cycling of IKK (active, inactive, and neutral IKK) induced by TNF.

This non-linear activation of IKK can easily arise from the trimerization of TNF receptors or complex oligomerization occurring in the early signaling cascade between the TNF receptor and activation of IKK (Chan, 2007; Hayden and Ghosh, 2014). Additionally, this simple modification from the minimal model enabled the digital activation of NF-κB observed in our previous study (Figure S2 and S3) (Tay et al., 2010). When we simulated the NF-κB behavior with this non-linear activation, we found that the resulting NF-κB dynamics corresponds better with the experimental observations from previous study and the current study. Further details about similarities or differences between the linear and non-linear activation are discussed in Figure S2 and S3. Also, to more accurately compare the simulation to the experimental results, we included other essential components in the signaling pathway that were omitted in the minimal model: the upstream inhibitor, A20, and the IκB kinases, IKK. IKK in neutral state is activated upon stimulation by TNF. The active IKK degrades IκBα initiating the nuclear translocation of NF-κB. The active IKK becomes inactive with help of A20, which then recovers back to the neutral state over time (Ashall et al., 2009; Tay et al., 2010). Hence, the cycling of IKK is controlled TNF and A20, and regulates the NF-κB dynamics.

We used object-oriented programming in C++ to conduct the simulations. The program generated text files containing relevant data and that we analyzed in MATLAB. Each grid point corresponds to a cell (programmed as an object) with individual concentrations of the proteins described in the model including the amount of nuclear NF-κB. We described the concentrations by ODEs except TNF, which is a PDE and solved using a two-dimensional FTCS method on a hexagonal grid, which is more precise than the traditional quadratic lattice. We used a fourth-order Runge-Kutta algorithm to solve the ODEs.

For the stochastic simulation with different activation threshold, the cell-to-cell variability was achieved through noise in the activation threshold of IKK, *C*_*TNF*_. Different thresholds were picked from a normal distribution with a mean value, *µ*_*CTNF*_, and standard deviation, *σ*_*CTNF*_.

The following table lists our modified equations and variables:

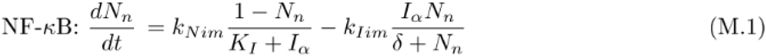

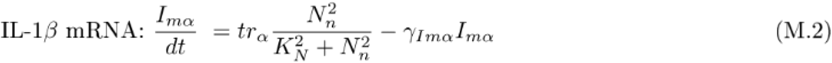

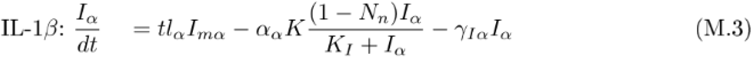

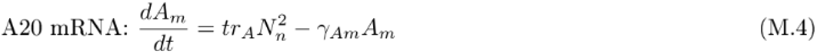

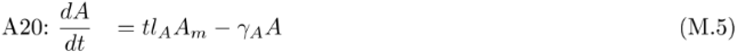

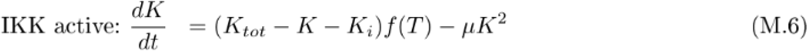

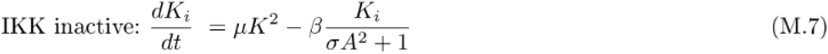

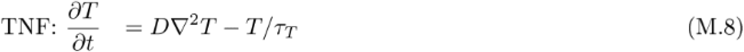

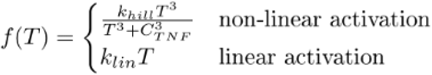

#### Parameters for the model

Many of the parameters used in our model originates from the measurement in the previous study (Hoffmann et al., 2002), while the rests are fitted to the experimental observations. The previous minimal model minimized the number of parameters by rescaling the variables (Krishna et al., 2006). However, we have only rescaled two parameters (NF-κB and TNF), and adapted the measured parameters as much as possible thereby increasing the credibility of the simulation.

### List of Parameters

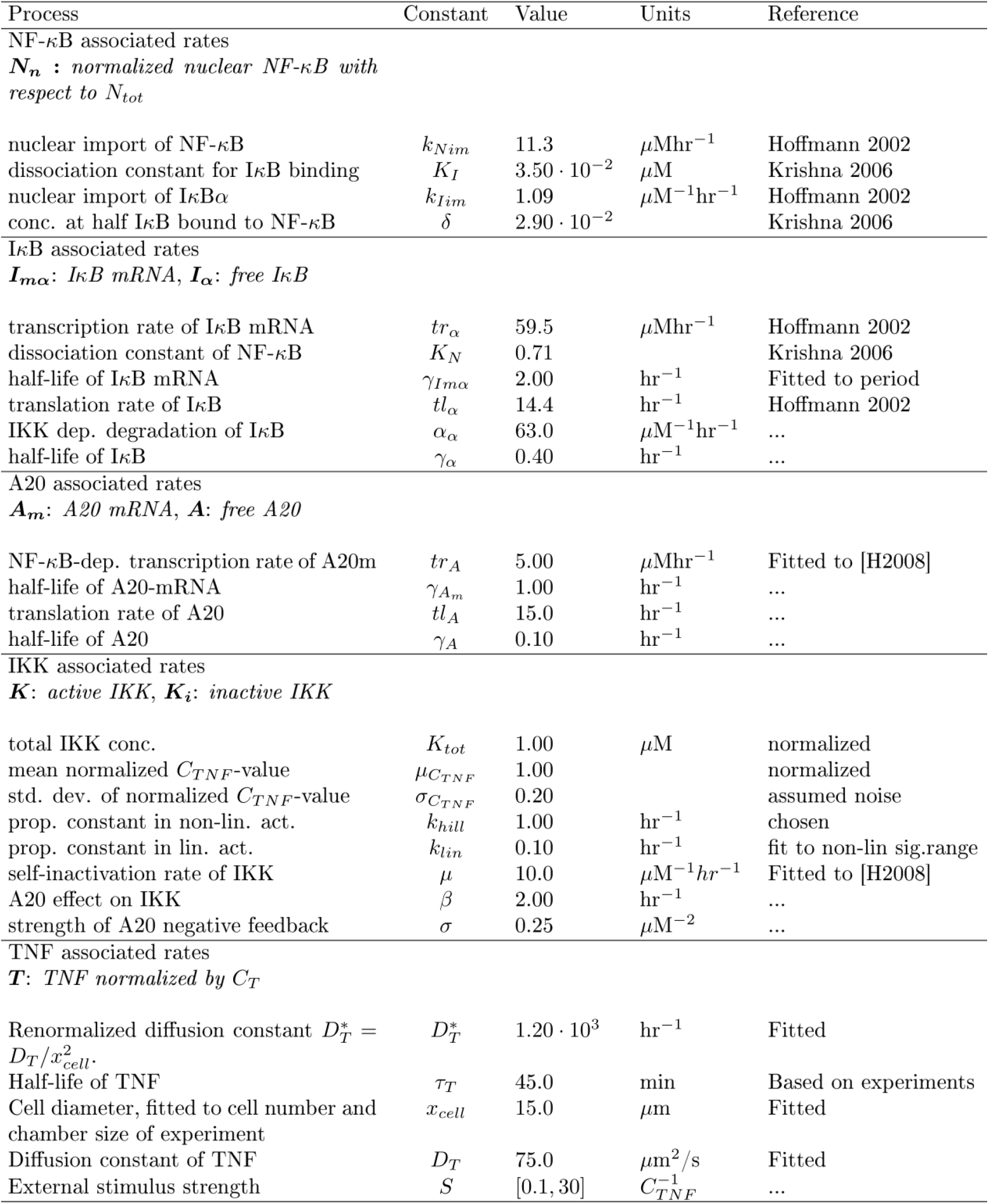

Parameters associated with A20 and IKK were fitted to reproduce qualitative findings in the previous study (Werner et al., 2008). In the previous study, the IKK response to a TNF stimulus peaked roughly 15 min after TNF stimulation and reached a steady state after roughly 30 min. The steady state level is determined by A20 where the active IKK level decreases with increasing A20. The mRNA level of A20 peaks around 30 min post stimulus and affects IKK dynamics after 45 – 60 min, see Figure S2. The unknown parameters in our model were fitted to reproduce these results. Other than unknown parameters, we increased the degradation constant, *γ*_*Imα*_, by two-fold. This change decreased the period between oscillations, which more closely matched the period observed in our experiments.

### TNF diffusion simulation

We simulated the diffusion explicitly on a grid with the similar boundary conditions as in our microfluidic experiments, where each grid point represents a single cell. Our two independent tests showed that the secretion from 3T3 fibroblast does not affect the neighboring cell’s NF-κB response (Figure S2 and S5). Hence, we assumed no secretion or stimulation between the neighboring fibroblasts. The diffusion parameter, *D*, was manually fitted, to reproduce the propagation of the first peak in the chamber, see Figure S3G – H.

For simulations of the test chamber, we used a lattice of size 16 × 130 cells ∼ 250 µm × (400 µm + 1300 µm) with three reflective boundaries and an open boundary at the top of the chamber. The cells where prepared in their steady states, which we found by running single-cell simulations for 100 h without external stimulus. At time t = 0, TNF stimulus was applied by setting the TNF concentration in the prechamber to a fixed level, S, (Figure 1B), which happened in a single time-step, dt. TNF then diffused through the chamber, similar to opening the experimental valve which separates receiver cells and the TNF-filled prechamber. For tissue simulations, the lattice consisted of 100 × 100 cells ∼ 1500 µm x 1300 µm and all boundaries were open. The stimulus was introduced at the center at time t = 0 and was maintained at the concentration S for a duration τ. This way, we could simulate the secretion of cytokine from a point source, such as a single macrophage. To simplify the secretion profile and to study the change in the tissue response depending on the characteristics (dose and duration) of the stimulus, we modeled the stimulus as a step-function.

In the case of co-culture experiment with both fibroblasts and a macrophage, we used the same chamber lattice, except we modified a single cell in the center of the pre-chamber to secret TNF (like a macrophage). At each time-step, the TNF concentration at the source, was adjusted such that the total secretion fits the experimentally measured secretion profiles from the macrophages (Figure S1) (Junkin et al., 2016). The secretion rates were normalized such that the macrophages with the strongest secretion activate fibroblast cells in the whole diffusion chamber.

### Downstream expression simulation

For the simulation for the downstream gene expression, we constructed two genes transcribed by NF-κB: an early gene which is expressed and decays fast and a late gene which is slowly accumulated. These two genes are described by two simple differential equations:

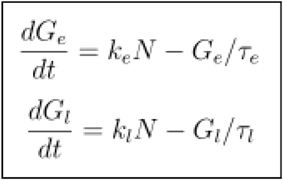

where the parameters are *k*_*e*_ = 4 h^-1^, *τ*_*e*_ = 0.25 h, *k*_*l*_ = 0.5 h^-1^, and *τ*_*l*_ = 16 h. With these simple changes in the expression characteristics, we demonstrated that different duration of the TNF stimulus can give a rise to regions with differentiated gene expression (Figure 6A).

