## Supplemental Figures for "Spatiotemporal NF-κB dynamics encodes the position, amplitude and duration of local immune inputs"

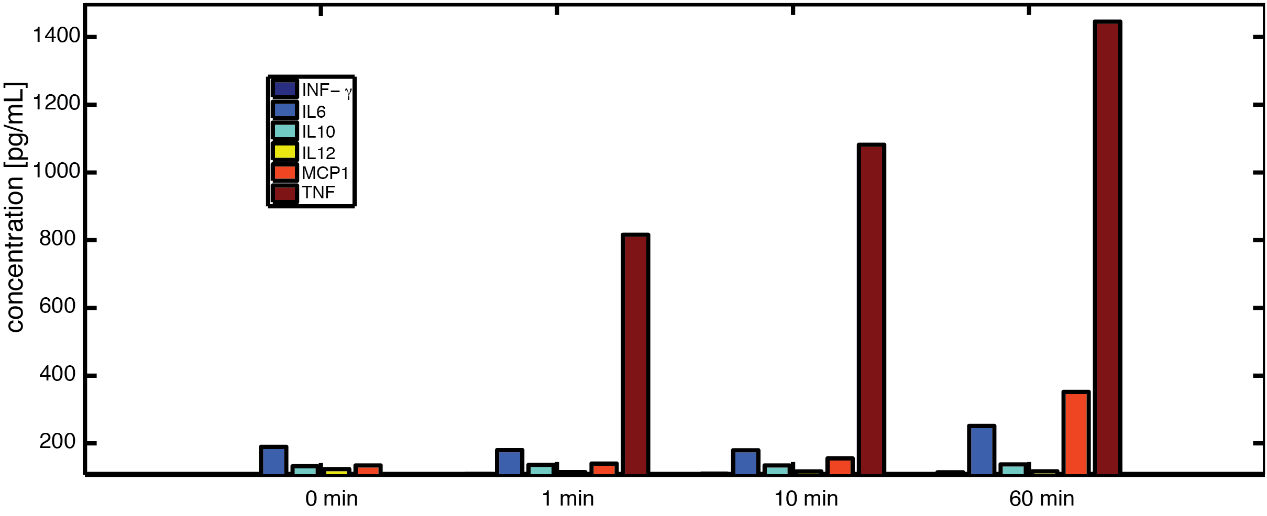

Figure S1: Secreted cytokines from RAW macrophages were measured with ELISA. Macrophage cells were cultured in a 12-well plate and stimulated with LPS (50 ng/mL) for 0,1,10 and 60 minutes. After stimulation, cells were washed three times, incubated further in fresh medium for 90 min, then the supernatant was collected for cytokine measurement. Using a mouse inflammatory cytokine kit, we measured six cytokines (INF-γ, IL-6, IL-10, IL-12, MCP1, and TNF) with antibody-coated beads, which was evaluated using FACS. Our results indicate that macrophages primarily secrete TNF in response to LPS stimulus.

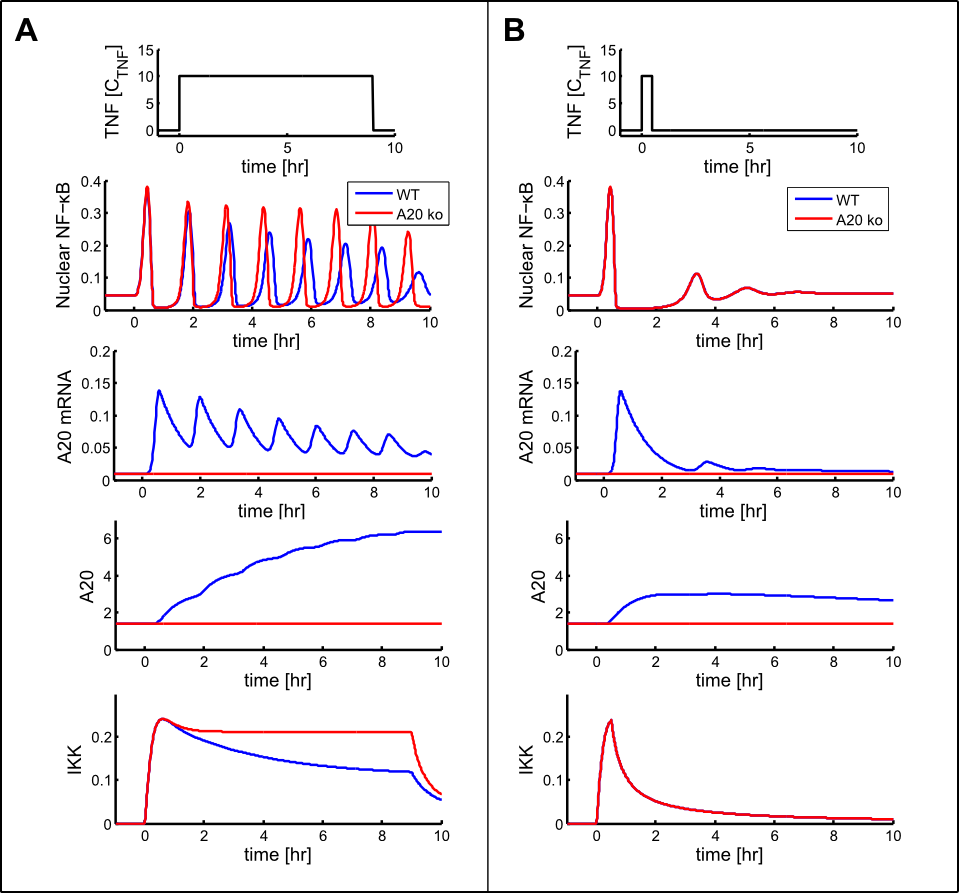

Figure S2: Comparison of WT and A20 knock-out simulations for short and long stimulus. NF-κB network was stimulated with 9 h pulse (A) and a 30 min pulse (B). For each input pulse, the dynamics of NF-κB translocation, A20 mRNA, A20 protein, and active IKK are shown, which showed similar kinetics to the experimental observations in the previous study (Werner et al., 2008). Our simulation shows that A20 affects the steady state level of active IKK, which in turn influences amplitude and period of the NF-κB translocation (A). However, for short stimulus (B), this effect is not observed due to fast decay of active IKK level after the stimulus ends.

**Linear versus non-linear stimulation:**

In our previous NF-κB model (Mengel et al., 2012), the activation of IKK through TNF was linearly correlated to the TNF level. However, another experimental study reported a constant area-under-curve (AUC) of the first peak over a wide range of TNF concentrations (Tay et al., 2010). To implement this in the model in the simplest way, we changed the activation of IKK from linear to a non-linear Hill-function (Eq. M9). Figure S3 shows how this change affects the characteristics of the response. Most notably, AUC of the first peak quickly saturates to a constant level in the non-linear model. In the linear model, AUC increases almost linearly as TNF level increases. Although this change slightly affected other response characteristics, the non-linear model successfully reproduced key findings in this study, such as number of oscillations or signaling distance.

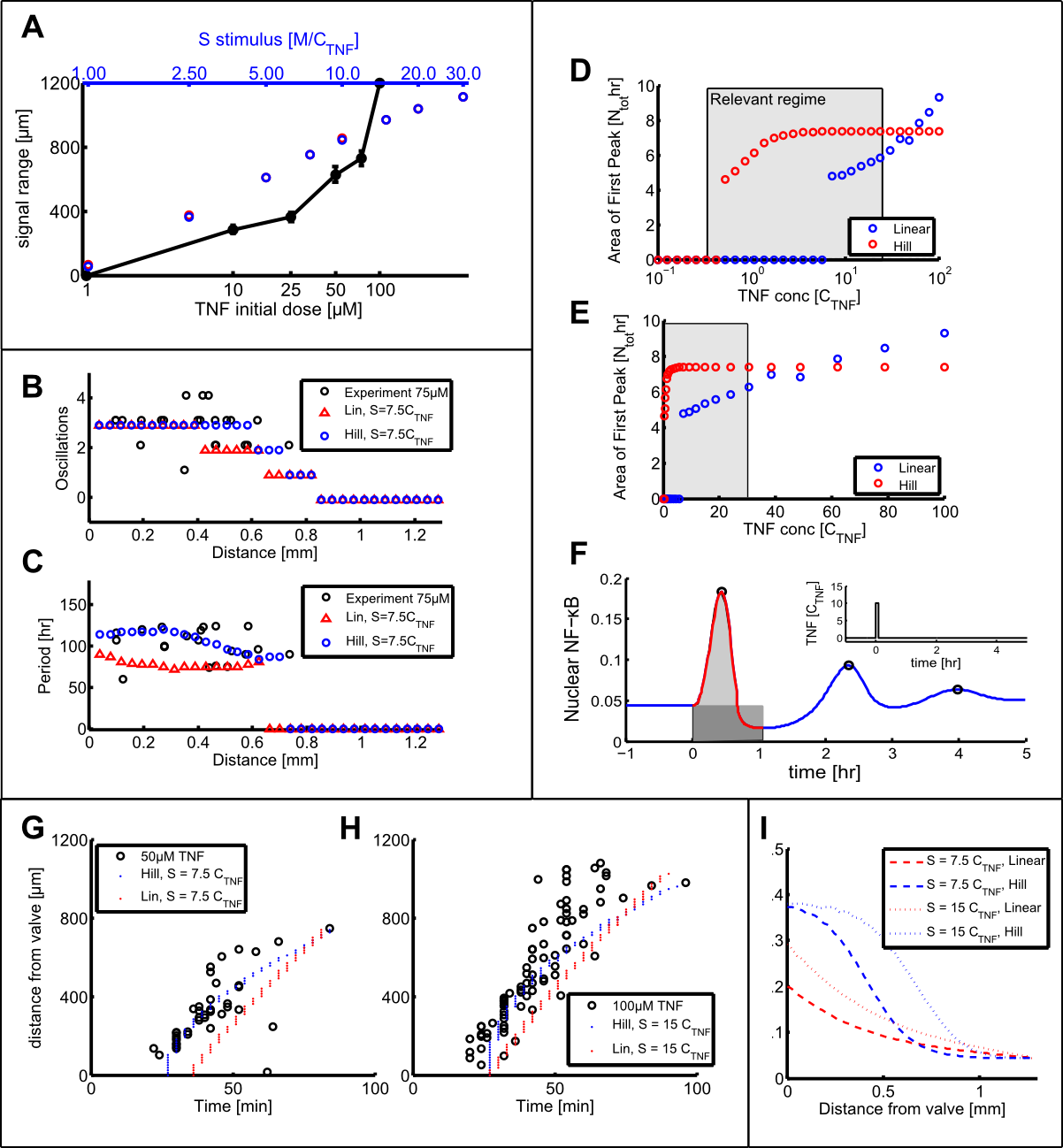

Figure S3: Comparison between linear and non-linear activation of IKK through TNF. In each simulation the chamber was exposed to a TNF-stimulus of the given strength and duration of 0.1 h (a stimulus pulse). No noise was included in the simulations to emphasize the average behavior of each model. The constant *k_lin_* is fitted so the signal range is similar for both models. The number of oscillations and the period is similar for both model types and neither is superior in describing the experimental data. The oscillation distribution in the chamber is steeper in the case of non-linear activation. The most remarkable difference in the two model types is observed in the NF-κB amplitude (I) and the area of the first peak (D+E). The area is calculated based on single cell simulations with no noise. For linear activation the area increases linearly with stimulus concentration while in the non-linear model the area quickly saturates to a constant area. (F) Subfigure of how the area under the first peak is calculated. The area is the gray area under the red curve minus the background activation, the rectangular box under the curve. The position of the first peak in time and space, (G+H) can qualitatively be described both by the linear and non-linear model.

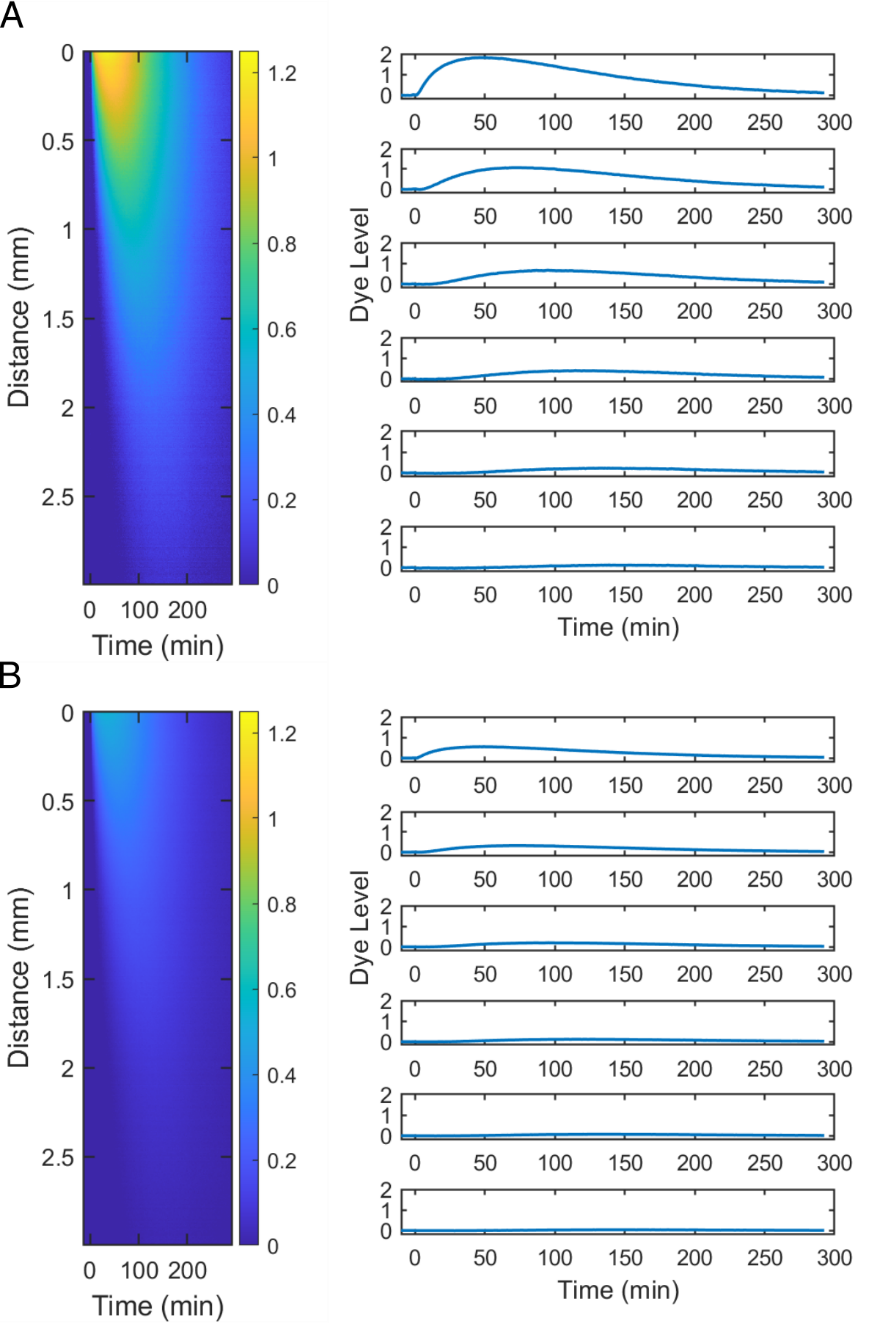

Figure S4: TNF dynamics at various distances from the source. (A) To evaluate the spatiotemporal dynamics of the diffusing TNF, high concentration of fluorescent dye was loaded in the source chamber, then the separating valve was opened at time t = 0. The fluorescence image was collected every 2 min for entire chamber, and plotted as heatmaps. The color of heatmaps corresponds to the logarithm of the fluorescence intensity. The time-course fluorescence change at six different distances (0.25, 0.75, 1.25, 1.75, 2.25, and 2.75 mm) was also evaluated and plotted on the right side. Here the dye level is in linear scale. (B) Similarly, the dye diffusion experiment was repeated with three tenth (3/10) concentration of what was used in (A). Hence, plots in (A) correspond to the 100 ng/ml source dose experiment, while plots in (B) correspond to the 30 ng/ml experiment.

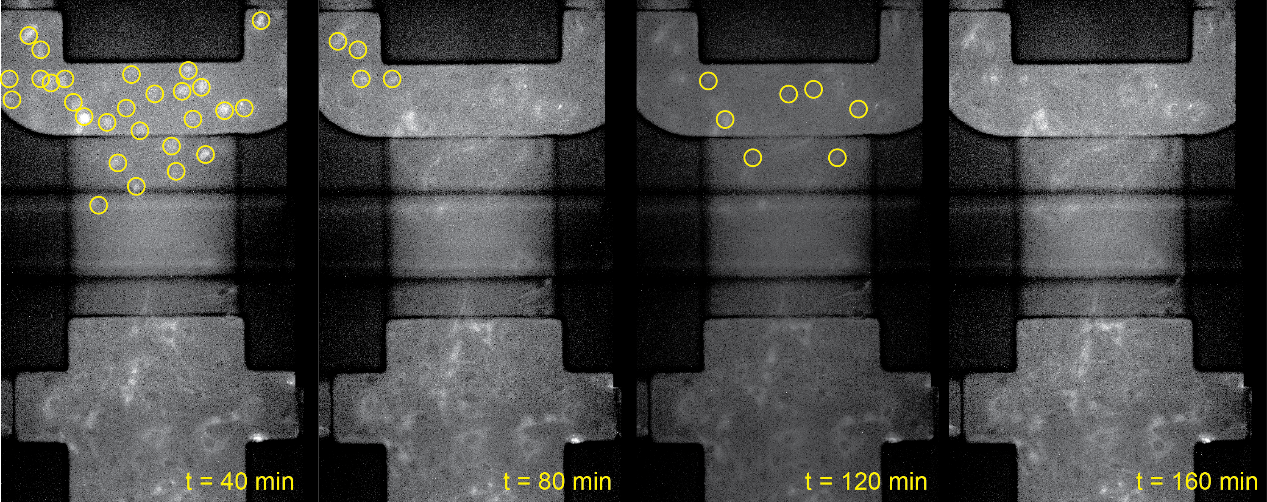

Figure S5: Test showing minimal cytokine secretion from fibroblast. Fibroblasts were loaded in both source and diffusion chamber. Fibroblasts in the source chamber were stimulated with 100 ng/ml TNF for 10 minutes, washed for 1 min with fresh medium, then the separating valve was opened to allow any secretion from the cells in the source chamber to diffuse to bottom. The figures show the fluorescence images highlighting the translocation of NF-κB. Yellow circles show the activated cells. Most cells in the source chamber were activated, while no cell became activated in the diffusion chamber even after 3 h.

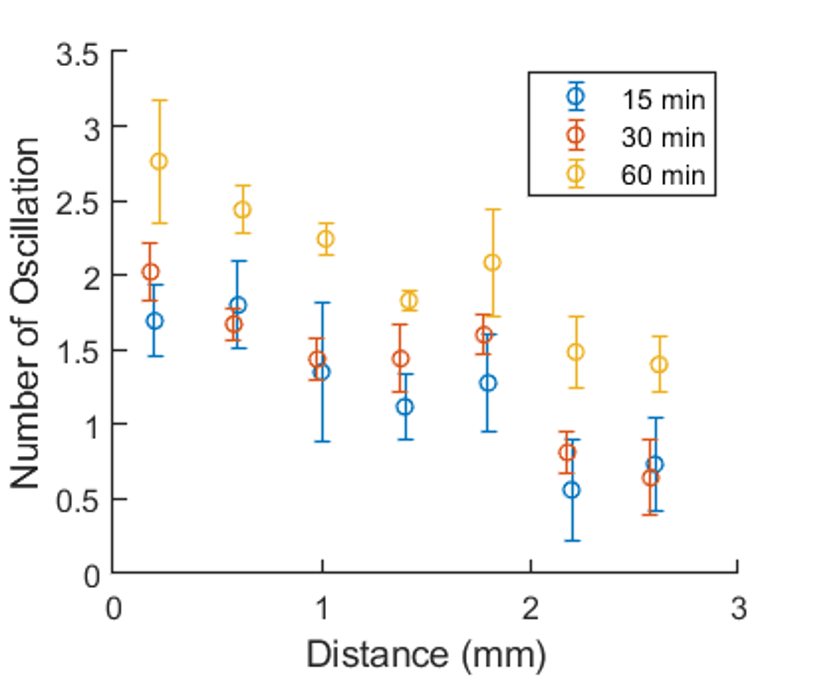

Figure S6. The longer cytokine secretion from a source results in the larger number of NF-κB oscillations at all locations. For each experiment with different secretion duration (Figure 3B), the cell chamber was divided into seven different regions, and the mean number of oscillations in each region was estimated. The results are displayed as mean ± SD from three replicates.

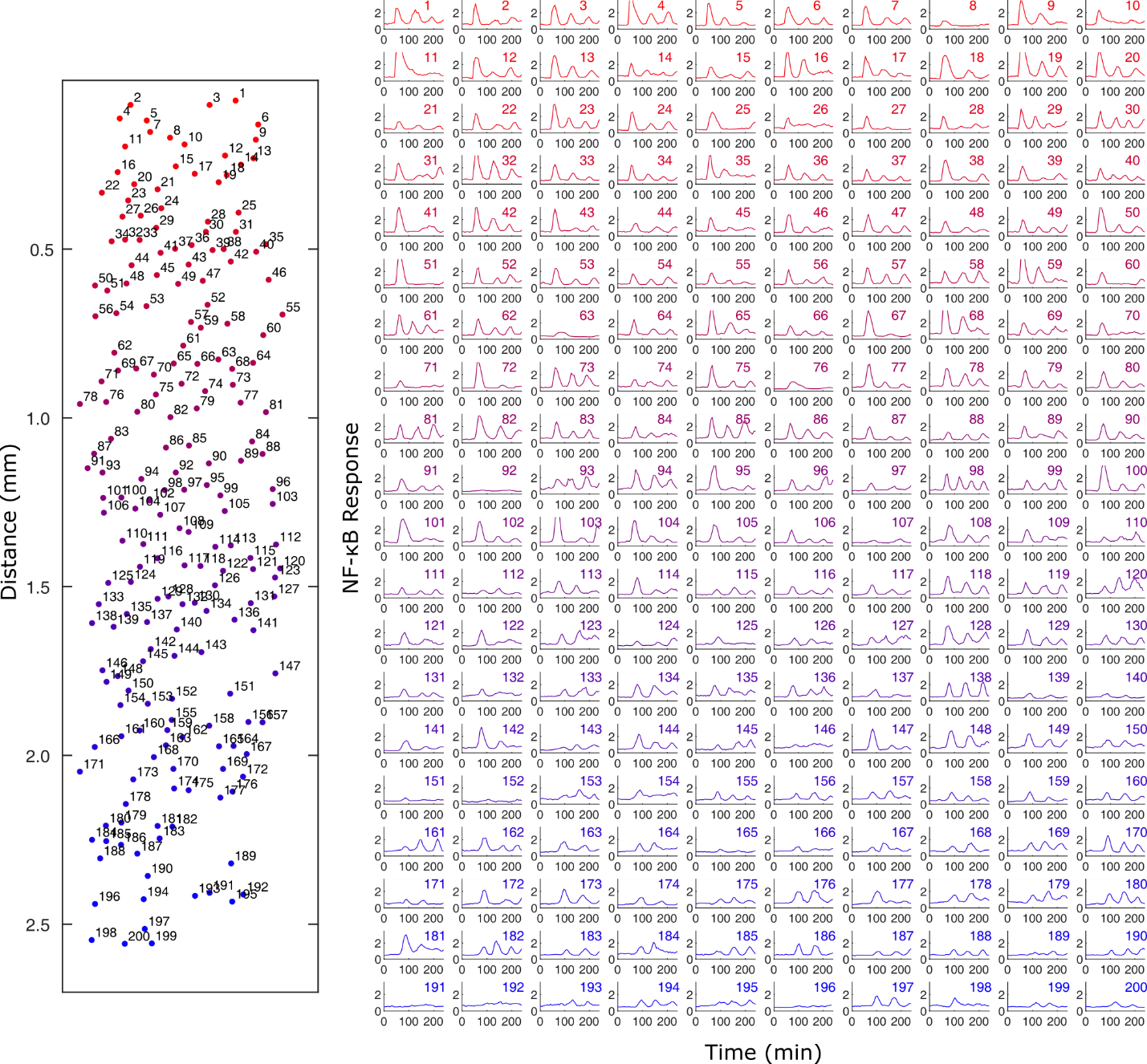

Figure S7: Spatially sorted single cell NF-κB dynamics for 100 ng/ml TNF source. 100 ng/ml TNF was loaded in the source chamber and the separating valve was opened at time t = 0. TNF diffused to the cell chamber and induced diverse NF-κB response based on the distance to the source.

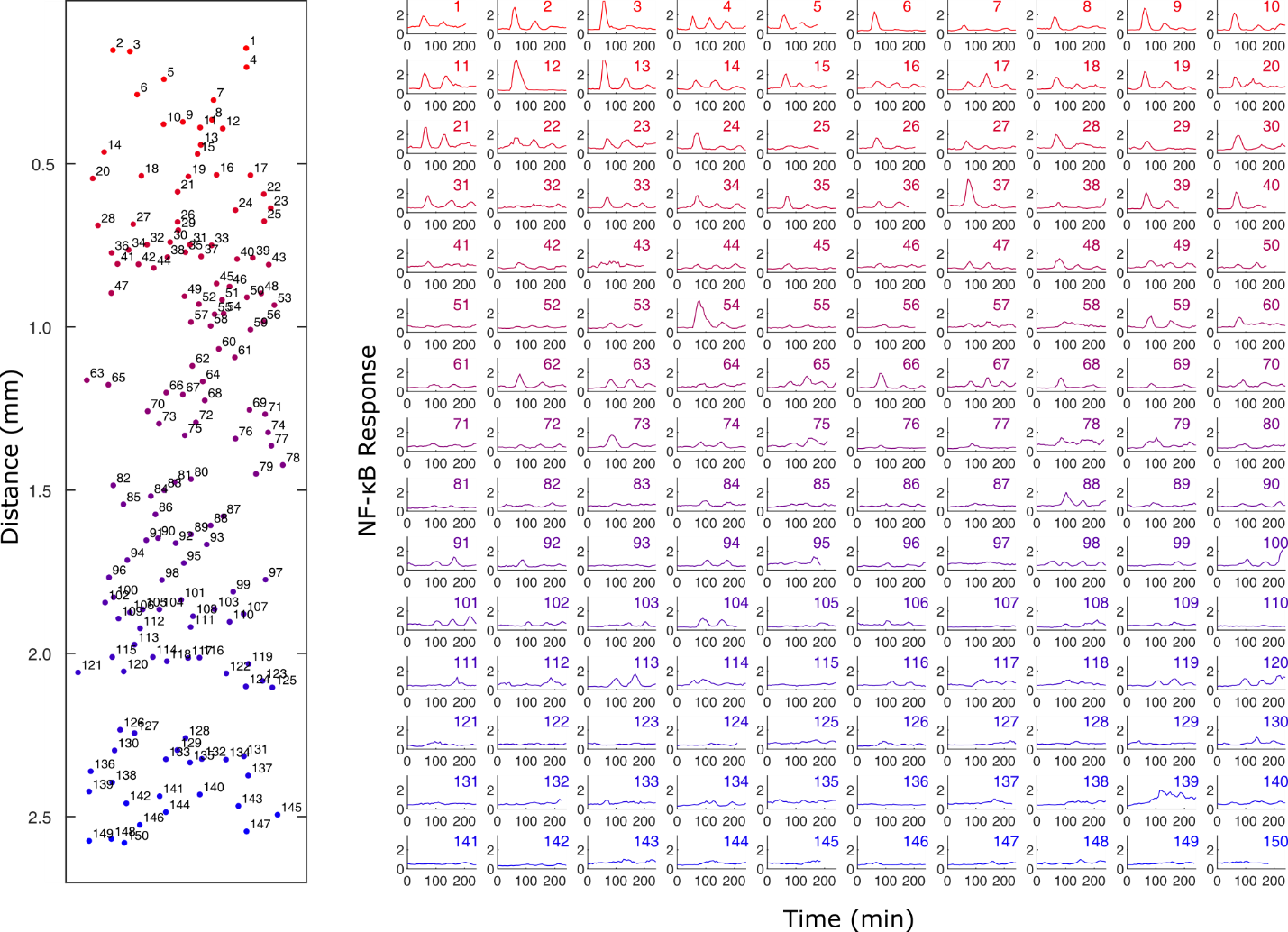

Figure S8: Spatially sorted single cell NF-κB dynamics for 30 ng/ml TNF source. 30 ng/ml TNF was loaded in the source chamber and the separating valve was opened at time t = 0. TNF diffused to the cell chamber and induced diverse NF-κB response based on the distance to the source.

**Data analysis:**

The NF-κB-response to a stimulus can be characterized by response amplitude, period, and number of oscillations. We specifically looked into how these characteristics change depending on the dose and distance from the localized TNF source. In the data analysis, an oscillation is defined as a peak in the active NF-κB-concentration, which is at least 25 % higher than the minimum value between the peak and the preceding and subsequent peak. If there is no other peak, we use the minimum value before and after the peak. Furthermore, the peak must be higher than the steady state level of nuclear NF-κB. This threshold is chosen to evaluate the oscillations, which are significantly different from the background noise. The response characteristics from the model and experiments are compared in Figure S9 ─ 10 for different concentrations of TNF. To match the chamber dimension used for the simulation, experimental results with the same chamber dimensions were used. We did not notice any significant difference in the results with smaller chamber, except the signaling range was downscaled proportional to the change in the dimension. The area under the first peak is calculated as the integral from the stimulus onset to the minimum after the first peak. To subtract the basal level, the steady state level of NF-κB multiplied by the duration of each peak is subtracted from the integral, see Figure S3F. The comparison between the model and experiment is shown in Figure S9 ─ 10.

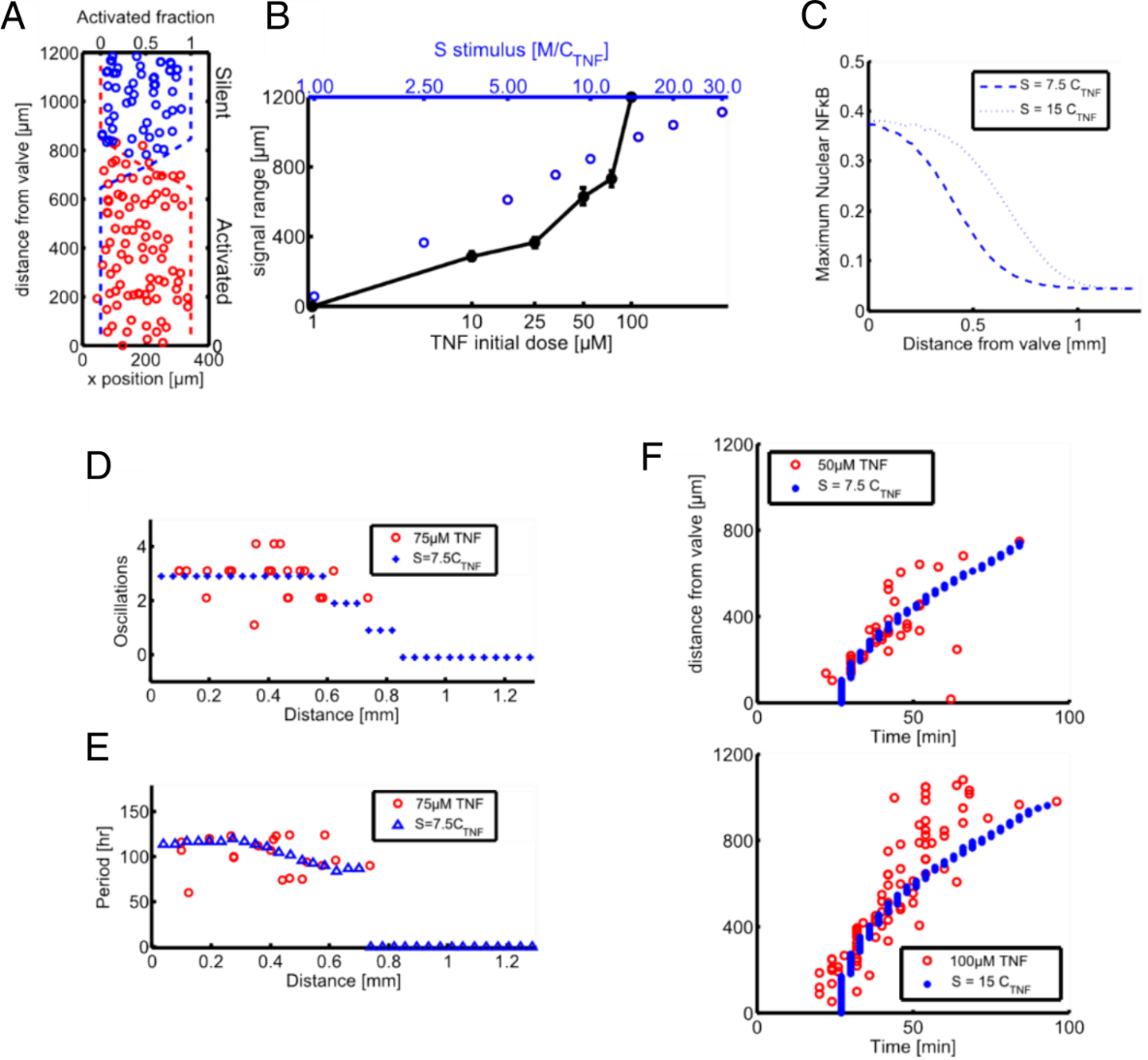

Figure S9: Detailed analysis of spatiotemporal NF-κB response induced by a local TNF source. (A) The diagram illustrates activation of cells in space. Every circle represents the position of a cell. Blue indicates inactive (silent) cells. Red represents activated cells. The dashed lines show the fraction of activated cells with respect to the distance from the valve. (B) The signaling range is dependent on local initial dose. We report the distance of how far the signaling propagates until the fraction of activated cells drops below 0.5. The black line shows the experimental results while blue reports the simulated values. (C) Predicted maximal nuclear NF-κB for two different doses of the TNF input. (D) Number of oscillations of single cells with respect to space. Red circles are number of oscillations from the experiments measured in single cells while blue dots are modeling results. (E) Period with respect to space is fairly constant. Red circles are experimentally measured periods of single cells while blue triangles represent the modeling output. (F) The diffusion constant of the model was refitted to the experimental spatiotemporal dynamics. The red circles represent the time and distance of the first activation.

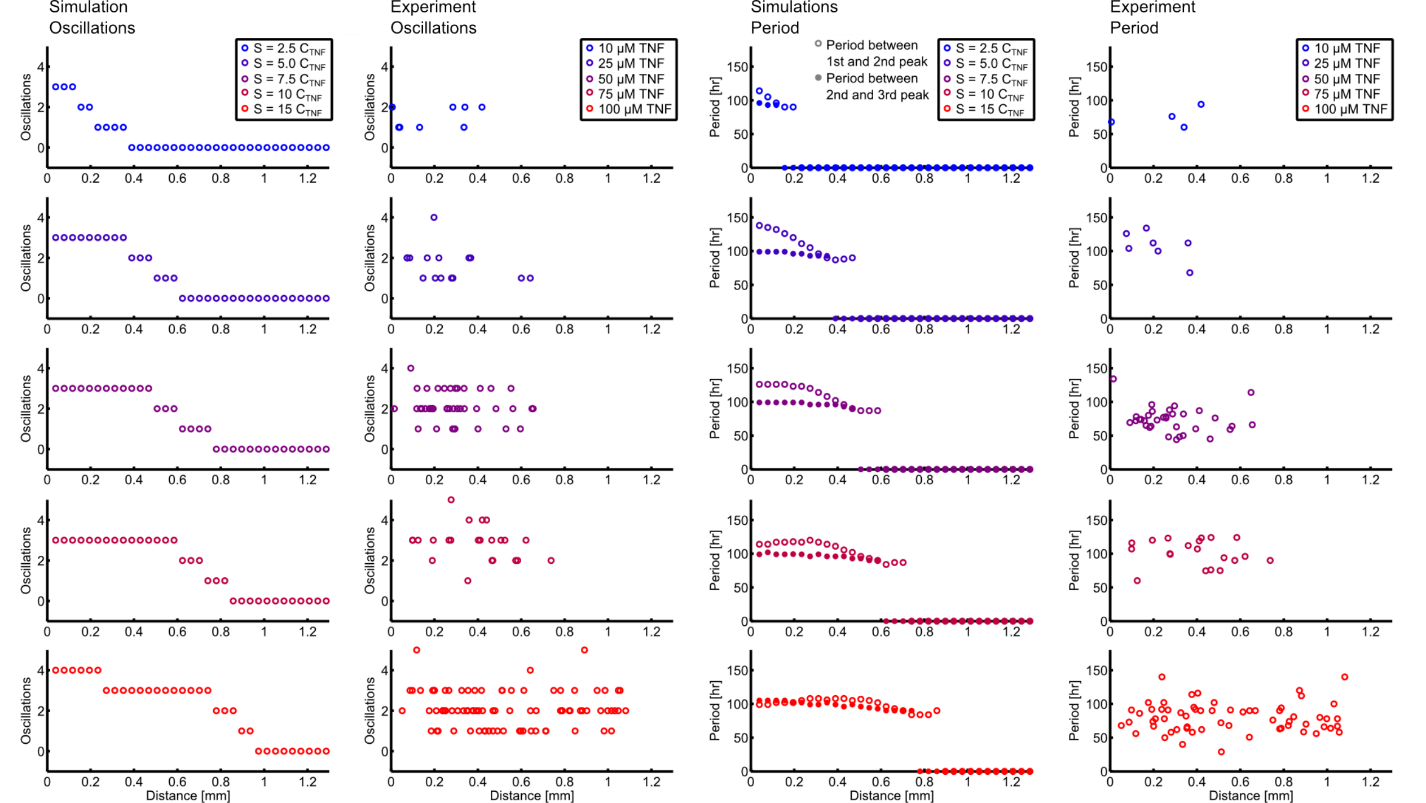

Figure S10: Dose dependency of oscillatory behavior. The plots show a screen of different local TNF initial conditions. Experimental and simulated results are reported. The left panels show number of oscillations while the right-hand side shows periods.

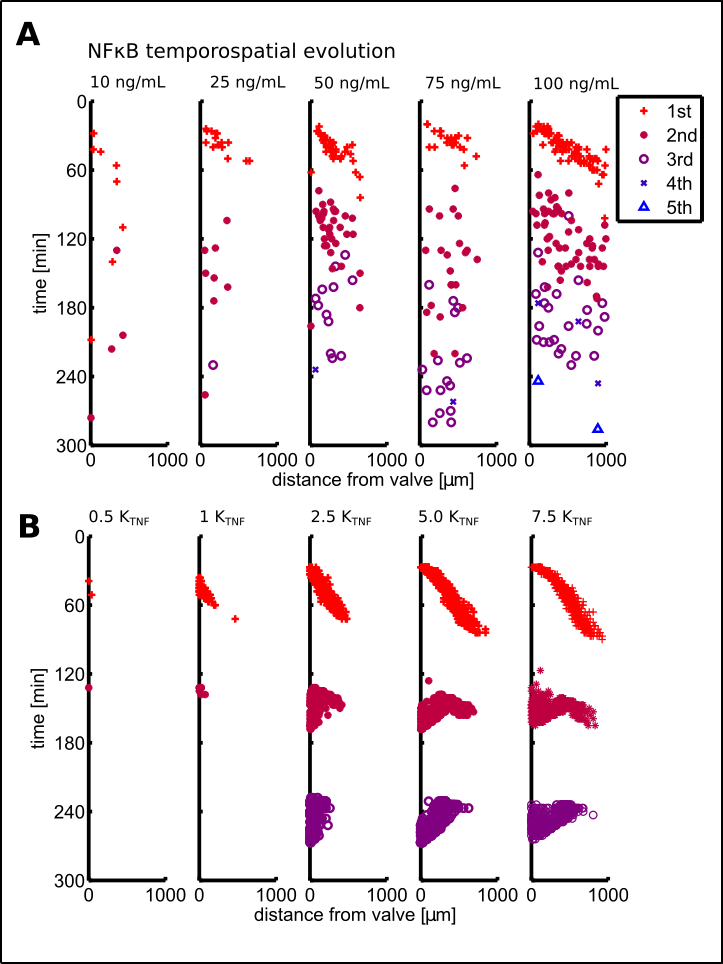

Figure S11: An initial wave of NF-κB activation is replaced by desynchronized oscillations. (A) The position of first to fifth oscillation observed in experiments. The number of oscillations increases with TNF concentration. The first oscillations are seen as a propagating wave in space while subsequent oscillations are more scattered space. (B) Simulation of NF-κB activation. The first response behaves as a travelling wave while following oscillations of NF-κB happens at similar times independent of space.

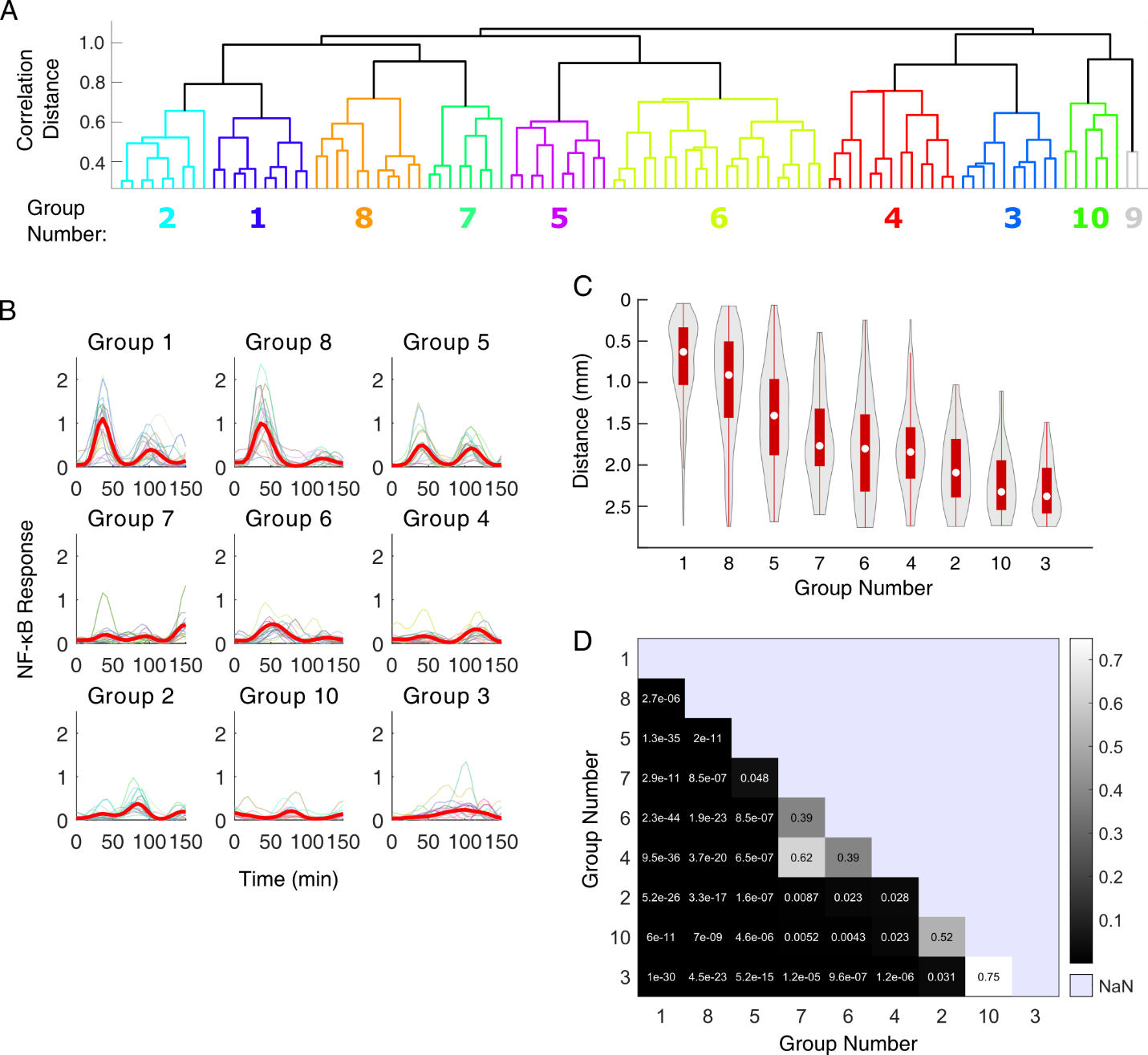

Figure S12: Cluster analysis of NF-κB activation in 30 ng/ml source TNF experiments showing distance dependency. (A) NF-κB responses in individual cells were clustered with ‘correlation’ method, and their relative distances were plotted. Group 9 contained few cells, and was considered as outlier. (B) Each subplot shows the mean (red line) and 20 examples of single cell NF-κB response in each group. The mean distance to the TNF source was calculated for cells in each group. The subplots were ordered based on the distance (Group 1 being closest, and Group 3 being farthest). (C) Violin- and box-plots of distance distribution in each cluster. (D) Kolmogorov-Smirnov statistics from a two-sample test comparing all combinations of distance distributions. Most pairs exhibited significant difference (< 0.05) except a few pairs which are positioned right next to each other.

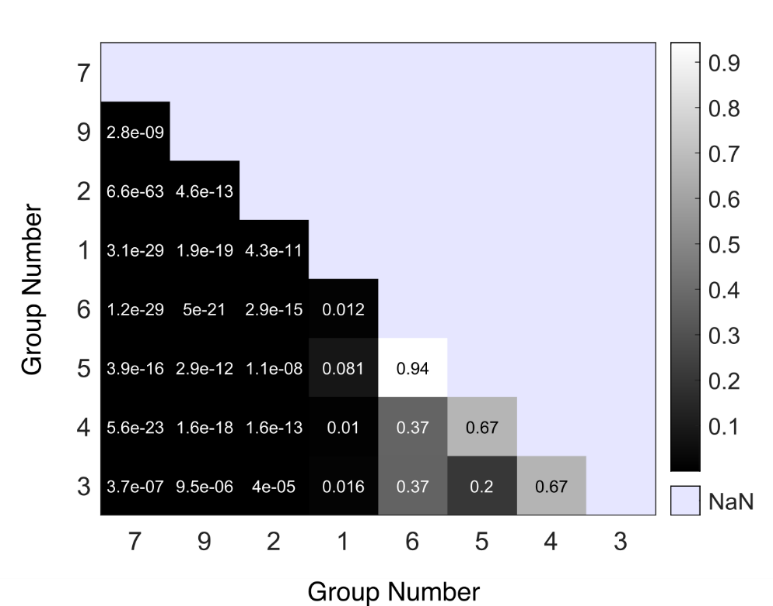

Figure S13: Kolmogorov-Smirnov (K-S) statistics for Figure 4C. Distance distribution from each group is compared with the distribution from another group using two-sample K-S. The distance distribution from Group 7, 9, 2, and 1 (in other words, groups closer to the cytokine source) showed significant difference (< 0.05) to all other groups, which indicates that the distinguishable NF-κB response behavior (Figure 4A − C) is linked to the cell’s distance to the cytokine source.

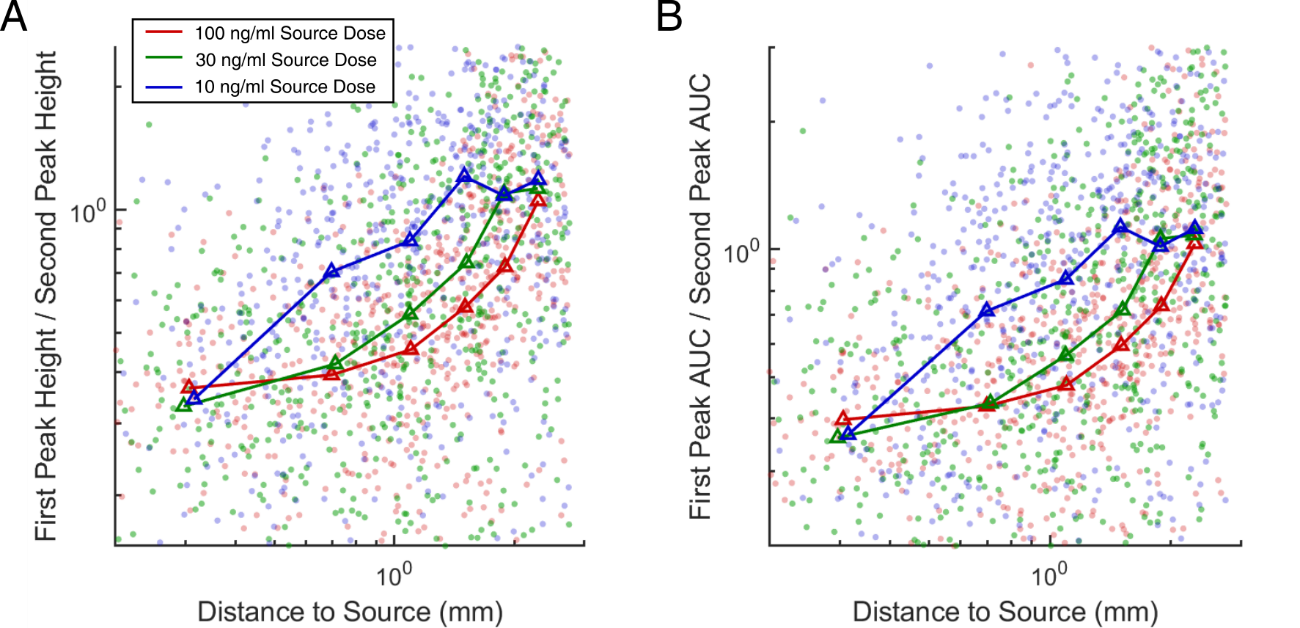

Figure S14. Ratio of the first to second peak vs. distance to the cytokine source. For various source doses, the ratio between the first and second peak amplitude (A) or area-under-curve (AUC) (B) is plotted against the distance to the source. Each dot in the scatter plots represents single cell data from 10 ng/ml (blue), 30 ng/ml (green), and 100 ng/ml (red) TNF source dose (700 cells were randomly chosen from each dose). Triangles indicate the mean of the single cell responses within 0.2 mm interval at six different distances from the source, 0.2 − 0.4, 0.6 − 0.8, 1.0 − 1.2, 1.4 − 1.6, 1.8 − 2.0, and 2.2 − 2.4 mm.

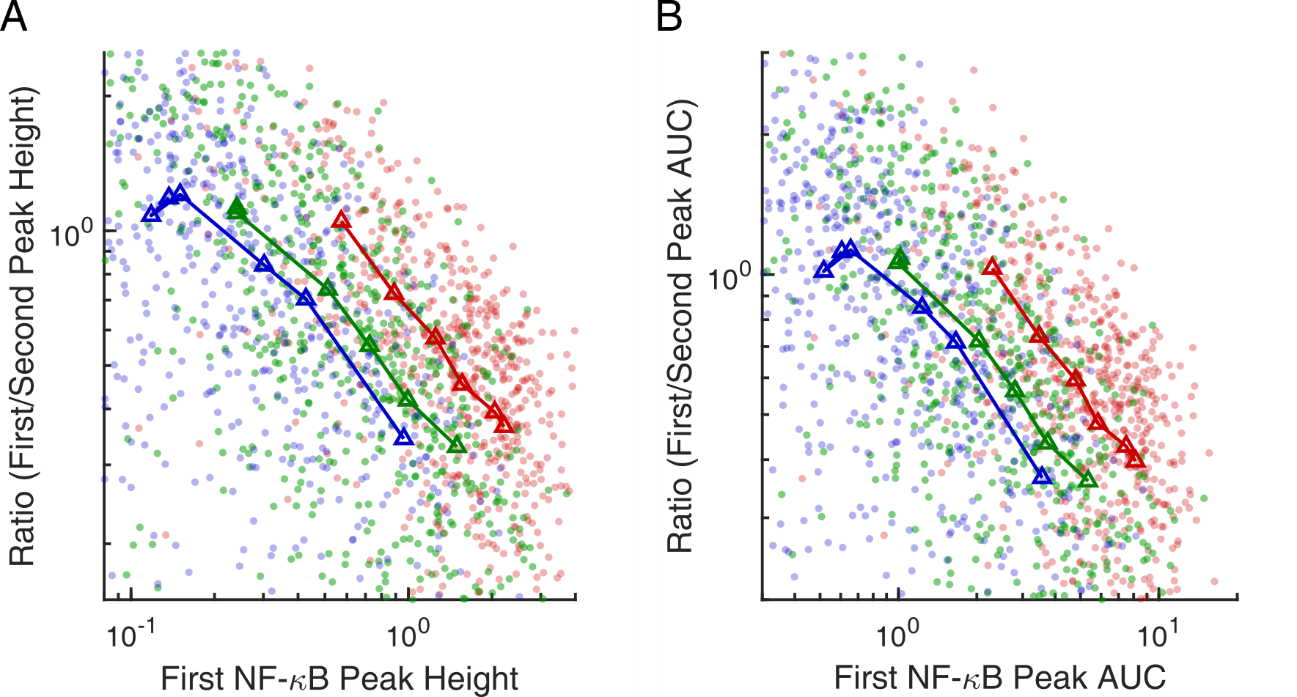

Figure S15: Ratio between the first and second peaks has information about the distance to the source cytokine. The ratio between the first and second peak magnitude (A) or area-under-curve (AUC) (B) was plotted against the first peak magnitude (A) or AUC (B). Each dot in the scatter plots represents single cell data from 10 ng/ml (blue), 30 ng/ml (green), and 100 ng/ml (red) TNF source dose (300 cells were randomly chosen from each dose). Triangles indicate the mean of the single cell response within 0.2 mm interval at six different distances from the source dose, 0.2 − 0.4, 0.6 − 0.8, 1.0 − 1.2, 1.4 − 1.6, 1.8 − 2.0, and 2.2 − 2.4 mm. Plots in (A) and (B) are largely similar, indicating both magnitude and AUC of the peak can be used to quantify the distance and dose of the TNF source.

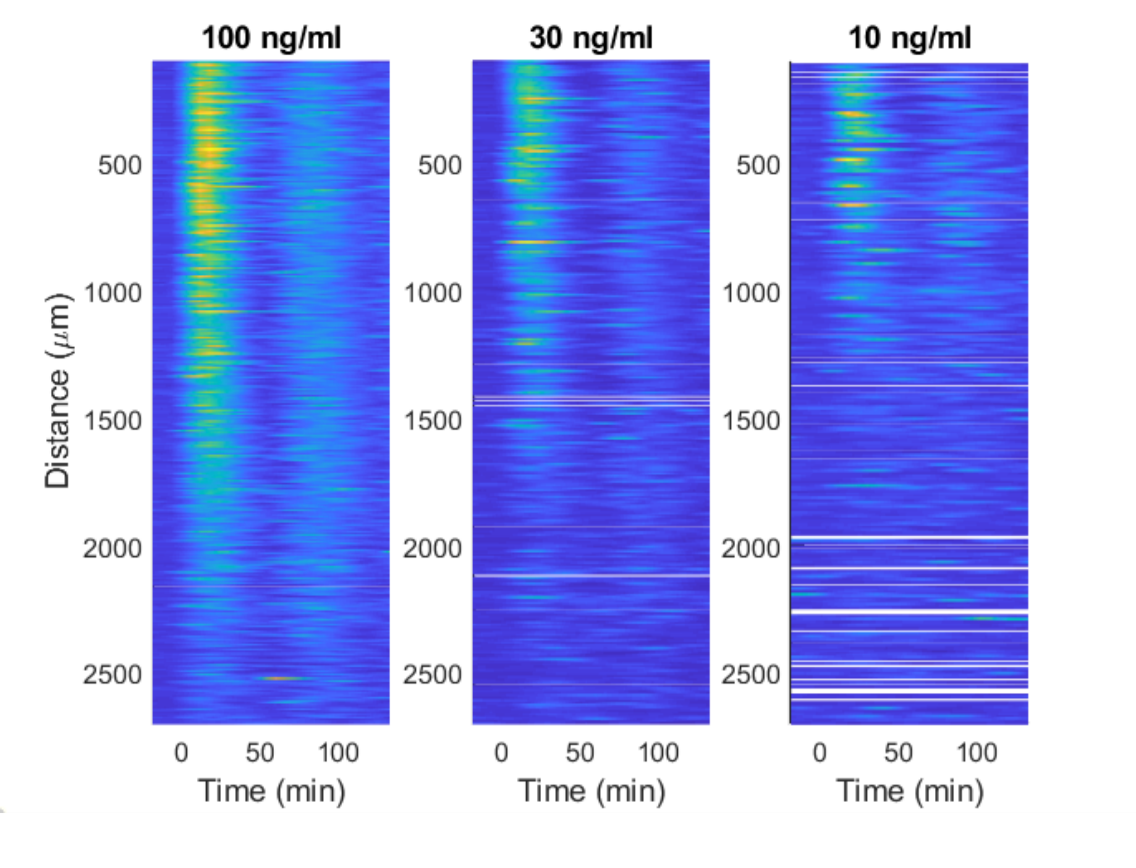

Figure S16: Onset timing of NF-κB activation was offset for accurate comparison of NF-κB response at different distance from different dose of source TNF.

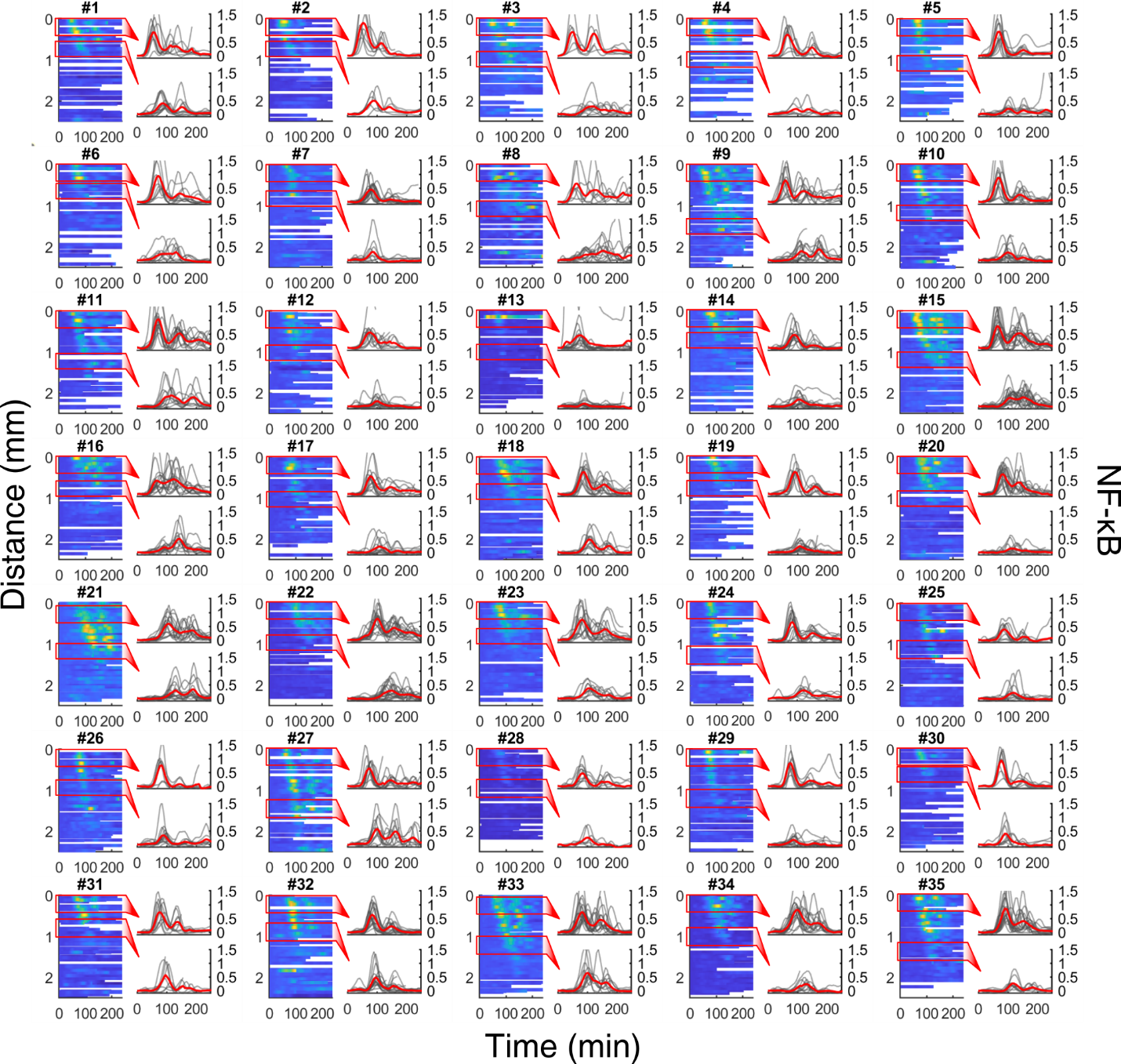

Figure S17: Co-culture experiments showing spatiotemporal NF-κB dynamics in fibroblast cells in response to the secretion from a single macrophage cell. A single macrophage cell was loaded in the source chamber, stimulated with 10 ng/ml LPS for 10 min, washed with fresh medium, then the separator was opened. The secretion from each cell was diffused to the cell chamber harboring fibroblasts, which activated the NF-κB translocation (heatmaps). For each sample, the response from fibroblasts at two different locations (red rectangles) is plotted in detail to study the difference in the NF-κB behavior based on the distance. The red line shows the mean of all response in each region.
